# The genome of *Caenorhabditis bovis*

**DOI:** 10.1101/766857

**Authors:** Lewis Stevens, Stefan Rooke, Laura C Falzon, Eunice M Machuka, Kelvin Momanyi, Maurice K Murungi, Samuel M Njoroge, Christian O Odinga, Allan Ogendo, Joseph Ogola, Eric M Fèvre, Mark Blaxter

**Author notes:** Author for correspondence: Lewis Stevens. Tree of Life, Wellcome Sanger Institute, Cambridge CB10 1SA, UK.

## Abstract

The free-living nematode *Caenorhabditis elegans* is a key laboratory model for metazoan biology. *C. elegans* is also used as a model for parasitic nematodes despite being only distantly related to most parasitic species. All ∼65 *Caenorhabditis* species currently in culture are free-living with most having been isolated from decaying plant or fungal matter. *Caenorhabditis bovis* is a particularly unusual species, having been isolated several times from the inflamed ears of Zebu cattle in Eastern Africa where it is believed to be the cause of bovine parasitic otitis. *C. bovis* is therefore of particular interest to researchers interested in the evolution of nematode parasitism and in *Caenorhabditis* diversity. However, as *C. bovis* is not in laboratory culture, it remains little studied and details of its prevalence, role in bovine parasitic otitis and relationships to other *Caenorhabditis* species are scarce. Here, by sampling livestock markets and slaughterhouses in Western Kenya, we successfully reisolate *C. bovis* from the ear of adult female Zebu. We sequence the genome of *C. bovis* using the Oxford Nanopore MinION platform in a nearby field laboratory and use the data to generate a chromosome-scale draft genome sequence. We exploit this draft genome to reconstruct the phylogenetic relationships of *C. bovis* to other *Caenorhabditis* species and reveal the changes in genome size and content that have occurred during its evolution. We also identify expansions in several gene families that have been implicated in parasitism in other nematode species, including those associated with resistance to antihelminthic drugs. The high-quality draft genome and our analyses thereof represent a significant advancement in our understanding of this unusual *Caenorhabditis* species.

## Introduction

The free-living nematode *Caenorhabditis elegans* is used extensively as a model for animal development, genetics and neurobiology. As the most well-studied species within the phylum Nematoda, *C. elegans* has also become a model for this extremely abundant and diverse group of animals, many of which are parasites (Blaxter *et al*. 1998; Blaxter & Koutsovoulos 2015). Attempts to understand the evolutionary origins and genetic basis of nematode parasitism often involve comparisons between parasitic nematode species and *C. elegans* (Bürglin et al. 1998; Gilleard 2004). However, *C. elegans* is only distantly related to most parasitic species which limits the efficacy of comparative studies (Blaxter & Koutsovoulos 2015). Recent years have seen significant progress in our understanding of *Caenorhabditis* diversity, with over 30 new species discovered in the last decade (Kiontke *et al*. 2011; Félix *et al*. 2014; Ferrari *et al*. 2017; Stevens *et al*. 2019). However, all of the ∼65 species currently in culture are free-living, with the vast majority having been isolated from rotting fruits and flowers (Kiontke *et al*. 2011; Félix *et al*. 2014; Ferrari *et al*. 2017; Stevens *et al*. 2019).

*Caenorhabditis bovis* (Kreis 1964) is therefore particularly unusual for a *Caenorhabditis* species, having been isolated several times from the outer auditory canal of Zebu cattle in Eastern Africa (Kiontke & Sudhaus 2006) and recently from Gyr cattle in South America (Cardona *et al*. 2010). *C. bovis* is believed to be the causative agent of bovine parasitic otitis, a disease which causes a range of symptoms including inflammation, dark brown discharge from the affected ear, and dullness (Msolla *et al*. 1993). In severe cases, bovine parasitic otitis can result in mortality (Msolla *et al*. 1993). As is typical for a *Caenorhabditis* species, *C. bovis* is believed to have a phoretic association with an invertebrate, with larvae of the Old World screwworm fly (*Chrysomya bezziana*) also being found in the ears of Zebu cattle (Msolla *et al*. 1989, 1993). It is unclear to what extent bovine parasitic otitis is caused directly by *C. bovis* or by bacterial and/or fungal infections and therefore to what extent *C. bovis* can be considered a parasite. Despite this, its close association with a vertebrate means that *C. bovis* is of particular interest to researchers interested in the evolution of nematode parasitism and in *Caenorhabditis* diversity. However, as *C. bovis* is not in laboratory culture, it remains little studied.

Here, by sampling cattle at livestock markets and slaughterhouses in Western Kenya, we successfully reisolate *C. bovis* from the ear of an adult female Zebu. We sequence the genome of *C. bovis* in a nearby field laboratory using the Oxford Nanopore MinION platform and use the data to generate a high-quality, chromosome-scale draft genome sequence. We exploit this genome to determine the phylogenetic relationships of *C. bovis* to other species in the genus *Caenorhabditis,* including *C. elegans*, and reveal the changes in genome size and content that have occurred during its evolution. We also reveal specific expansions in several gene families which may play a role in its unusual lifestyle. The high-quality draft genome and the analyses presented here represent a major step forward in our understanding of this unusual and understudied *Caenorhabditis* species.

## Results

### Reisolation of *C. bovis*

We sampled a total of 44 cattle of various ages and breeds at livestock markets and slaughterhouses in three counties in Western Kenya (Fig. 1A; Table S1). Sampling was performed by washing the outer auditory canal of each animal with cotton wool soaked in physiological saline (Fig. 1B) which was subsequently inspected under a dissecting microscope. We identified only a single instance of bovine parasitic otitis. The affected animal was an adult female Zebu which was sampled at a livestock market in Chwele, Bungoma County (Fig. 1A). The animal is believed to have originated from West Pokot County (Fig. 1A). Although we noted no obvious clinical symptoms, the cotton wool sample had an unpleasant odour, consistent with previous reports of bovine parasitic otitis (Msolla *et al*. 1993). We isolated approximately 50 live nematode larvae from the sample which were subsequently cultured on *E. coli*-seeded agar plates. The cultures thrived at 37°C on both nematode growth medium (NGM) and horse blood agar plates. Adult nematodes were identified as members of the genus *Caenorhabditis* based on their morphology (presence of a prominent pharyngeal bulb and filiform female tail). The morphology of the adult male tail (anteriorly closed fan, ray pattern with gap between GP2 and GP3, and a bent gubernaculum) was consistent with previous descriptions of *C. bovis* (Kreis 1964; Sudhaus & Kiontke 1996).

**Figure 1:**
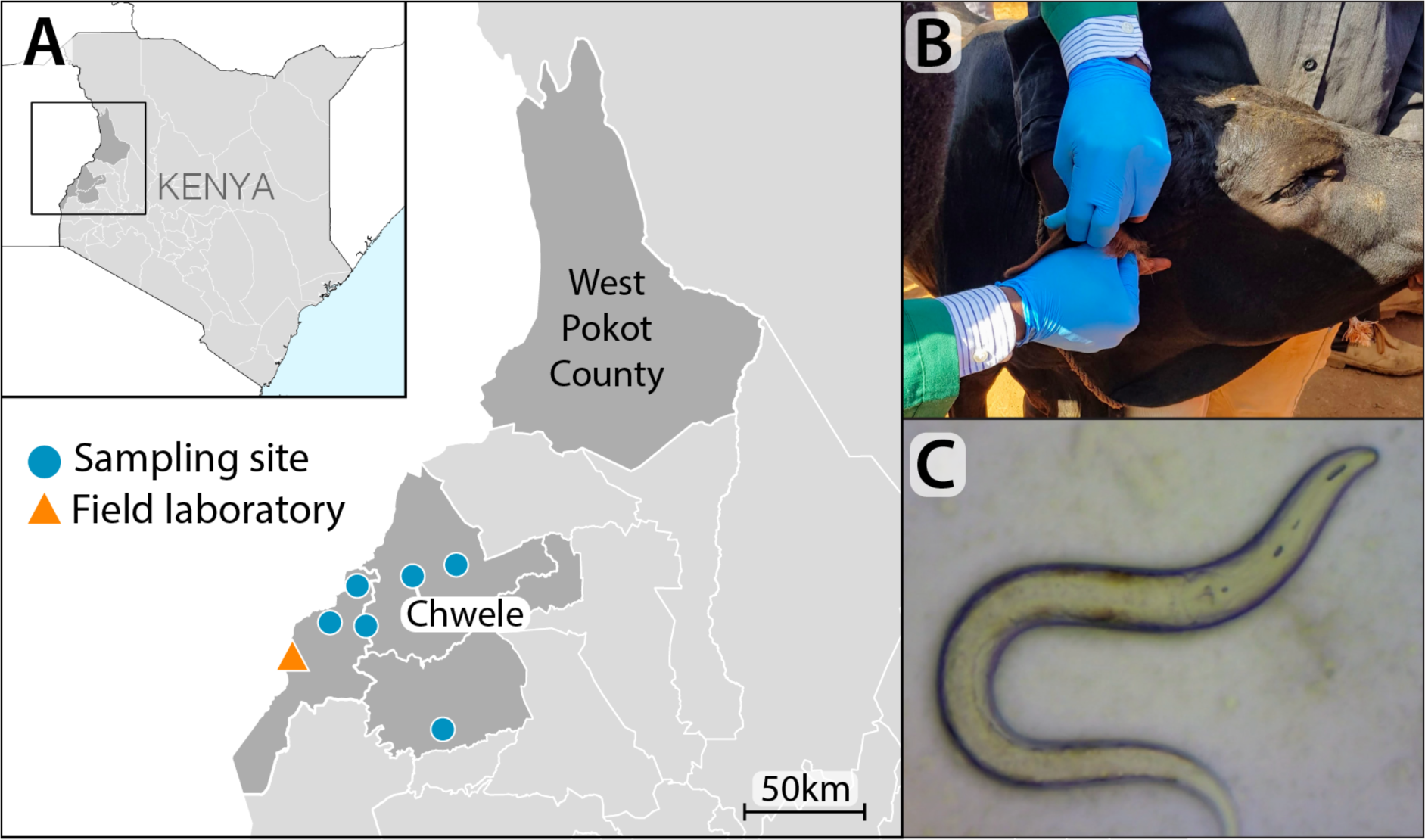
Cattle sampling and nematode isolation A: Sampling locations in Western Kenya. We isolated *C. bovis* from an adult female Zebu sampled at a livestock market in Chwele. The animal was believed to have originated from West Pokot County. The location of the field laboratory in Busia is also shown. GPS coordinates and the number of animals sampled at each site can be found in Table S1. **B:** An animal being sampled using cotton wool soaked in physiological saline. **C:** Adult female *C. bovis* under a stereo microscope (*C*. *bovis* adults are ∼1mm in length (Kreis 1964)).

**Table 1:**
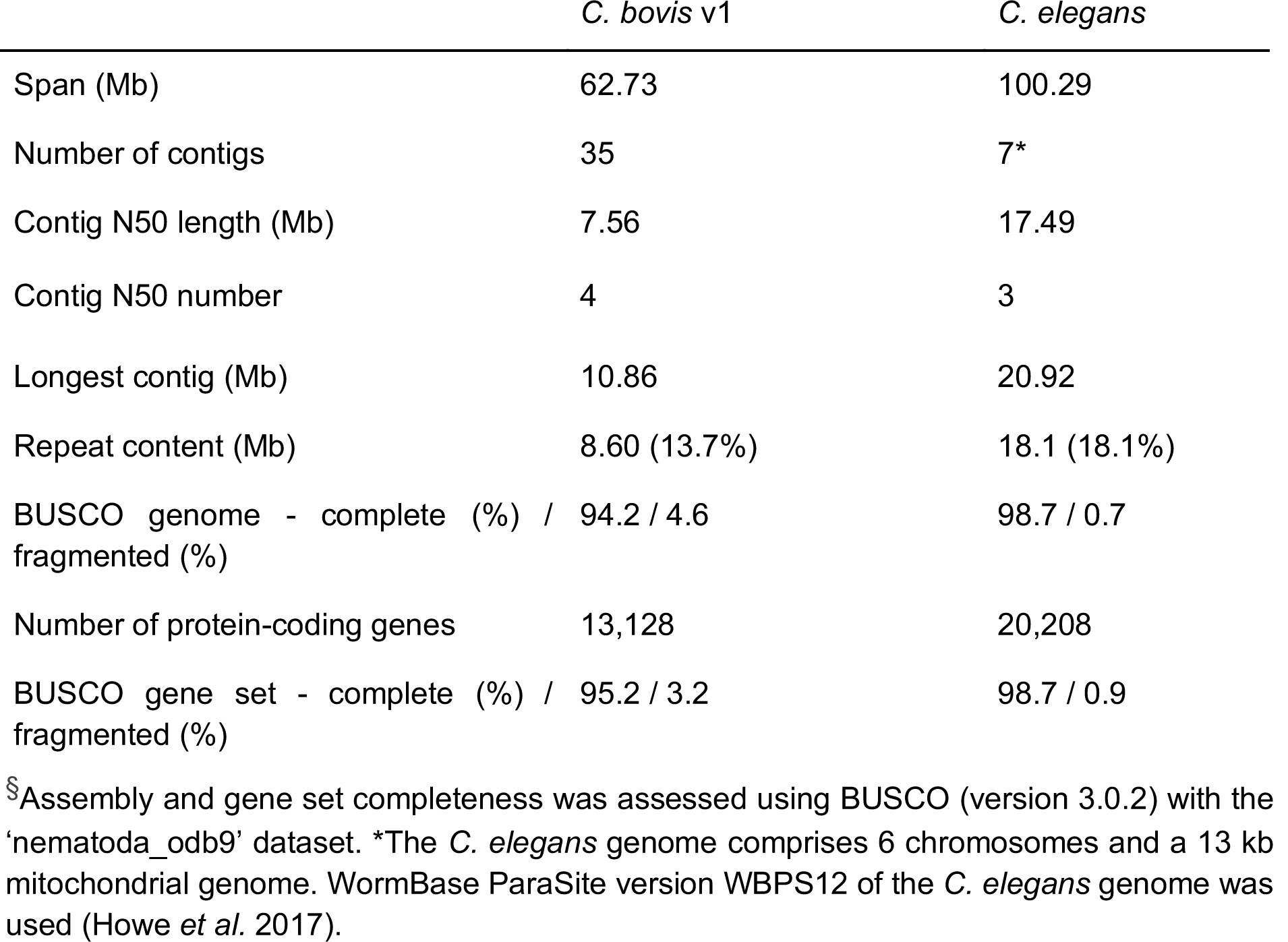
Genome and gene set metrics^§^ for Caenorhabditis bovis assembly v1.0

### A high-quality, chromosome-scale *C. bovis* reference genome

We sought to generate a high-quality reference genome for *C. bovis*. We took advantage of the portability of the Oxford Nanopore MinION platform and sequenced the genome of *C. bovis* in a field laboratory in Busia, Western Kenya (Fig. 1A). We generated 11.3 Gb of sequence data representing ∼180-fold coverage of the *C. bovis* genome using two MinION v9.4 flow cells. Read length N50s were 11.4 kb and 4.3 kb, respectively, with the longest read spanning 242 kb (Table S2; Fig. S1). We also sequenced the genome to ∼210-fold coverage (13.3 Gb) using the Illumina MiSeq platform at the BecA-ILRI Hub in Nairobi, Kenya. We identified and discarded reads originating from contaminant organisms, including several bacterial species which are known mammalian pathogens, using taxon-annotated GC-coverage plots (Fig. S2).

We assembled the *C. bovis* genome using the MinION long-reads and corrected residual sequencing errors in the assembly using the Illumina short-reads. The resulting assembly comprises 35 contigs spanning 62.7 Mb with a contig N50 of 7.6 Mb, with half of the assembly contained in just 4 contigs (Fig. 2A,B; Table 1). The assembly is highly complete, with 94.2% of a conserved set of nematode genes being present and fully assembled. The genome contains surprisingly little heterozygosity (0.13% as estimated using kmer spectra; Fig. S3). Using protein sequences predicted from the genomes of related nematodes as homology evidence, we predict 13,128 protein-coding genes in the *C. bovis* genome. We note that this number is considerably lower than the number of genes predicted in the genomes of other *Caenorhabditis* species (Stevens *et al*. 2019). However, the gene set contains 95.1% of a conserved set of nematode genes (Table 1), suggesting that the reduced count is not due to an incomplete gene set.

**Figure 2:**
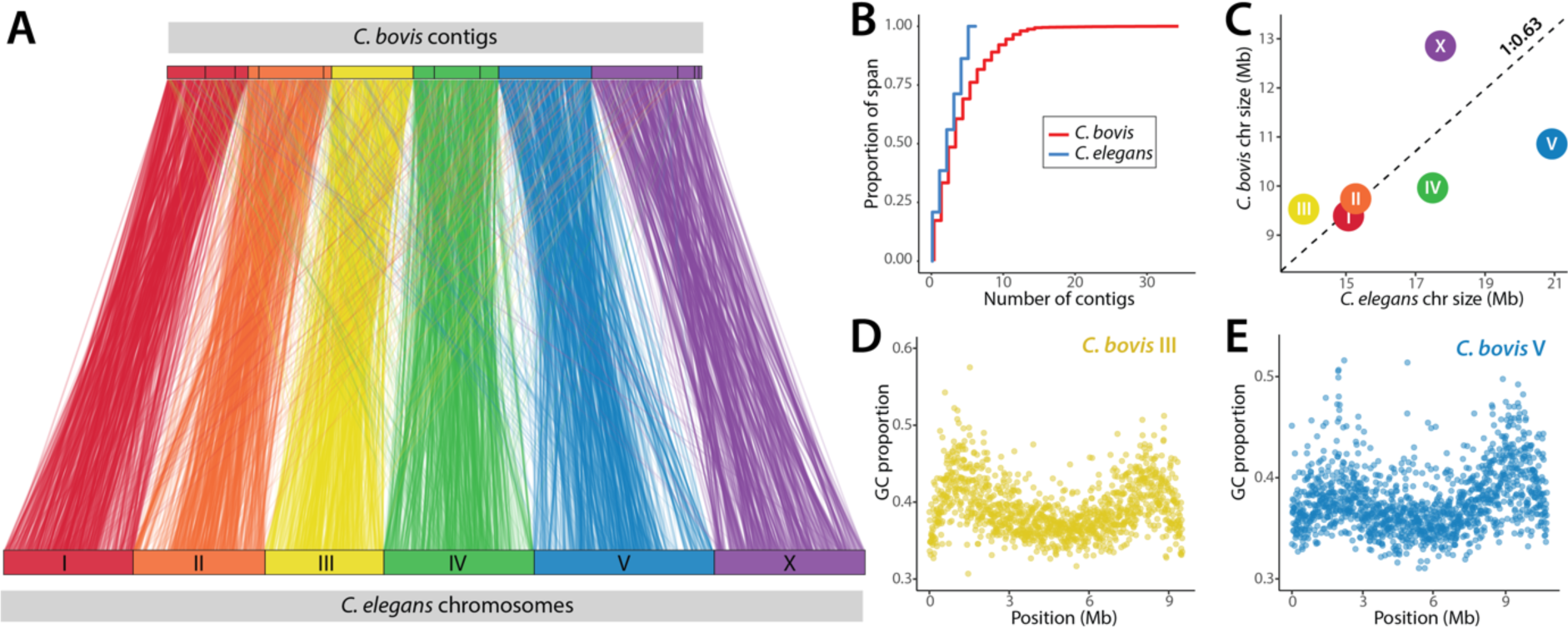
A high-quality, chromosome-scale *C. bovis* reference genome A: Highly conserved linkage groups enable the assignment of *C. bovis* contigs to the six *C. elegans* chromosomes. 15 contigs representing 99.4% of *C. bovis* assembly are shown. Lines represent the position of 7,706 orthologues between *C. bovis* and *C. elegans*. **B:** Cumulative length as a proportion of assembly span plot of *C. bovis* and *C. elegans*. **C:** Chromosome size in *C. bovis* and *C. elegans.* Dotted line represents the expected chromosome size based on the proportion of overall genome size between *C. elegans* and *C. bovis* (1:0.63). **D, E:** Patterns of variation in GC content (using an 8 kb sliding window) in *C. bovis* contigs 3 (chromosome III) and 1 (chromosome V), respectively, are consistent with the arms and centers organisation present in the chromosomes of other *Caenorhabditis* species.

Chromosomal linkage groups are highly conserved in *Caenorhabditis* (Hillier et al. 2007). We defined 7,706 orthologues between *C. bovis* and *C. elegans* and exploited this conservation to assign 15 contigs representing 99.4% of the *C. bovis* assembly to the six *C. elegans* chromosomes (Fig. 2A; Fig. S4). Chromosomes III and V are represented by single contigs, suggesting that these contigs represent complete *C. bovis* chromosomes. Both contigs also show patterns of variation in GC content characteristic of the arms and centres organisation present in the chromosomes of other *Caenorhabditis* species (*C. elegans* Sequencing Consortium 1998; Hillier *et al*. 2007; Yin *et al*. 2018) (Fig. 2D,E). The remaining chromosomes are each represented by 3-4 contigs (Fig. 2A; Fig. S4).

### The position of *C. bovis* within *Caenorhabditis*

We sought to reconstruct the phylogenetic relationships of *C. bovis* to other species in the genus *Caenorhabditis*. We clustered over a million protein sequences predicted from the genomes of 32 other *Caenorhabditis* species and two outgroup taxa, *Diploscapter coronatus* and *Diploscapter pachys*, into orthologous groups and selected 1,167 single-copy orthologues. Alignments of these orthologues were concatenated to form a supermatrix which was used to reconstruct the *Caenorhabditis* phylogeny using maximum likelihood. Our phylogenomic analysis resulted in a well-supported phylogeny (Fig. 3) which was largely congruent with previously published phylogenies (Kiontke *et al*. 2011; Slos *et al*. 2017; Stevens *et al*. 2019). We recover *C. bovis* as sister to *Caenorhabditis plicata* with maximal support (bootstrap value of 100). The clade containing *C. bovis* and *C. plicata* is early-diverging within the genus *Caenorhabditis*, and the branches separating *C. bovis* and *C. plicata* are long, indicating that *C. bovis* is highly diverged from all other sequenced species, including *C. elegans*.

**Figure 3:**
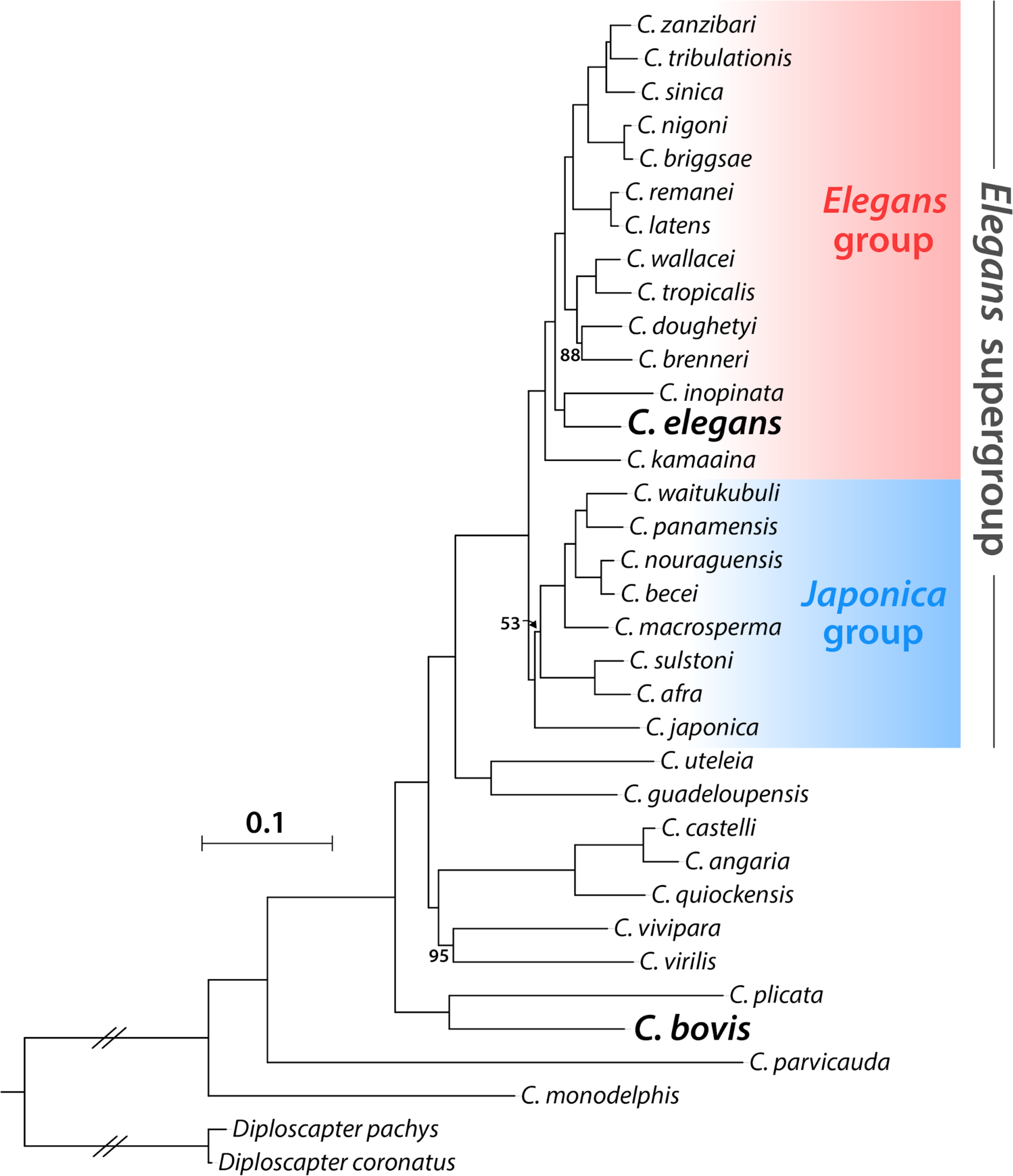
**The phylogenetic position of *C. bovis* within *Caenorhabditis*** Phylogeny inferred using a supermatrix of 1,167 single-copy orthologues under the general time reversible substitution model with gamma-distributed rate variation among sites (GTR + Γ). *C. bovis* and *C. elegans* are highlighted in bold. The tree is rooted with the two *Diploscapter* species. Branch lengths are in substitutions per site; scale is shown. Bootstraps were 100 unless noted as branch annotations. Major clades as defined by (Kiontke *et al*. 2011) are highlighted.

### Genome shrinkage in *C. bovis*

At 62.3 Mbp, the *C. bovis* genome is the smallest *Caenorhabditis* genome published thus far and nearly 40% smaller than the *C. elegans* genome (*C. elegans* Sequencing Consortium 1998; Stevens *et al*. 2019). The majority of the difference in span (22.1 Mbp or 58%) is due to differences in protein-coding gene content, with the *C. bovis* genome containing 7,080 fewer predicted genes than the *C. elegans* genome. The 13,128 *C. bovis* genes span 35.1 Mb (56% of the *C. bovis* genome) while the 20,208 *C. elegans* genes span 57.2 Mb (57% of the *C. elegans* genome). Several of the most highly reduced gene families in *C. bovis* comprise G protein-coupled receptors (GPCRs) (Table S3), with differences in the number of GPCRs alone accounting for 16% (1,139) of the difference in the number of genes between *C. bovis* and *C. elegans*. The *C. bovis* genome also contains considerably less repetitive DNA than the *C. elegans* genome (Table 1), with 12% of the *C. bovis* genome (7.4 Mb) comprising transposable elements compared with 14% of the *C. elegans* genome (13.9 Mb) (Table S4).

While all six *C. bovis* chromosomes are smaller than their homologous chromosomes in *C. elegans,* the differences are not equal (Fig. 2C). Chromosome V is most reduced, being 49% smaller than *C. elegans* chromosome V and containing less than half as many genes (2,302 and 4,992, respectively). Interestingly, the X chromosome is most well conserved in terms of size and gene content, with the *C. bovis* X chromosome being only 28% smaller than the *C. elegans* X chromosome and containing 20% fewer genes (2260 and 2782, respectively).

Despite the genome being considerably smaller, *C. bovis* genes contain more introns than their *C. elegans* orthologues (8.6 and 6.6 introns per gene, respectively; Table S5). This is consistent with previous analyses which have found that early-diverging *Caenorhabditis* species have retained more ancestral introns than their in-group relatives (Kiontke *et al*. 2004; Slos *et al*. 2017). However, *C. bovis* introns are, on average, less than half the size of *C. elegans* introns (157 bp and 319 bp, respectively; Table S5). Therefore, despite containing more introns, *C. bovis* genes contain on average less intronic DNA than their *C. elegans* orthologues (1270 bp and 2375 bp of intronic DNA per gene, respectively; Table S5).

### Expanded gene families in the reduced *C. bovis* genome

We sought to identify features of the *C. bovis* genome which may relate to its unusual ecology. Using the orthology clustering set described previously, we compared the *C. bovis* gene set to those of 32 other *Caenorhabditis* species. Despite having a substantially reduced number of genes, we identified several *C. bovis*-specific expansions in gene families which have independently been implicated in parasitism in other nematode species.

We find evidence for two duplications of the orthologue of the *C. elegans* gene *pgp-11* in *C. bovis,* resulting in three distinct copies (Fig. S5). All other *Caenorhabditis* species, except for *C. monodelphis,* possess a single orthologue of *pgp-11* (Fig. S5). In *C. elegans*, *pgp-11* encodes a P-glycoprotein which belongs to the ATP binding cassette (ABC) transporter family (Sheps *et al*. 2004). P-glycoproteins have been implicated in resistance to ivermectin and other antihelminthic drugs in several parasitic nematode species (Xu *et al*. 1998; Bourguinat *et al*. 2008; Bartley *et al*. 2009). *C. elegans* strains which lack *pgp-11* function show increased susceptibility to ivermectin (Janssen *et al*. 2013b) and genetic variation in the orthologue of *pgp-11* in the horse parasite *Parascaris equorum* is associated with decreased susceptibility to ivermectin (Janssen *et al*. 2013a).

The orthologue of the *C. elegans* gene *far-8*, which encodes a fatty acid and retinol (FAR) binding protein (Garofalo *et al*. 2003), has undergone two duplications in *C. bovis*, resulting in three distinct copies (Fig. S6). The secretion of FAR proteins by parasitic nematodes is thought to interfere with host immune defences and/or play a role in the acquisition of essential lipids (Bradley *et al*. 2001). A family of proteins containing Kunitz-type serine protease inhibitor domains has undergone expansion in *C. bovis* (Fig. S7). The majority of species, including *C. elegans,* possess a single member of this family, while *C. bovis* possess five. A Kunitz-type serine protease inhibitor secreted by the hookworm *Ancylostoma ceylanicum* has been shown to be capable of inhibiting mammalian host proteases (Milstone *et al*. 2000). In addition, a family of galectin-domain containing proteins appears to be restricted to *C. bovis.* Galectins are actively secreted by several parasitic nematode species and may interfere with mammalian host immune responses (Turner *et al*. 2008; Kim *et al*. 2010; Wang *et al*. 2014).

## Discussion

Here, we reisolated *C. bovis* from the ear of a female Zebu (*Bos taurus indicus*) in Western Kenya. We sequenced the genome of *C. bovis* using the Oxford Nanopore MinION platform in a nearby field laboratory and used the data to generate a high-quality, chromosome-scale reference genome. We exploited this genome sequence to reconstruct the phylogenetic relationships of *C. bovis* to other *Caenorhabditis* species, and identified expansions in gene families that may be associated with the unusual lifestyle of *C. bovis*.

The low level of heterozygosity in the *C. bovis* genome is surprising. Genomes of outcrossing *Caenorhabditis* species typically contain extremely high levels of polymorphism (Cutter *et al*. 2006; Dey *et al*. 2013) which can complicate genome assembly (Barrière *et al*. 2009). To circumvent these issues, *Caenorhabditis* species are often deliberately inbred over multiple generations (e.g. by sibling mating) prior to sequencing (Stevens *et al*. 2019). While it is likely that our *C. bovis* cultures underwent some population bottlenecking during isolation and the subsequent two-week period of laboratory culture, if *C. bovis* has similar levels of heterozygosity to other outcrossing *Caenorhabditis* species, this alone is not sufficient to explain the low levels of heterozygosity we observe. Instead, it seems that the *C. bovis* population we sampled from is naturally highly inbred, suggesting that a very small number of nematodes are transported between hosts and that gene flow between demes is extremely rare. Resequencing other isolates would allow us to test if this is true of all *C. bovis* populations.

The placement of *C. bovis* as sister to *C. plicata* is intriguing. *C. plicata* has been isolated from carrion (once from a dead elephant in Kenya and once from a dead pine marten in Germany) and has a phoretic association with carrion beetles (Volk 1950; Sudhaus 1974; Kiontke & Sudhaus 2006). *C. plicata* is therefore the only *Caenorhabditis* species currently in culture that has been found in association with a vertebrate, with all others having been isolated from rotting plant or fungal matter (Kiontke *et al*. 2011; Félix *et al*. 2014; Ferrari *et al*. 2017; Stevens *et al*. 2019); Marie-Anne Felix, Lise Frezal, Matthew Rockman, Christian Braendle, John Wang, Michael Ailion, Erik Andersen, Asher Cutter, pers. comm.). In recent years, worldwide sampling has led to the discovery of many new *Caenorhabditis* species, but all efforts have been focussed in habitats that resemble the decaying vegetable matter habitat identified as the home of *C. elegans* (Félix & Braendle 2010; Kiontke et al. 2011; Félix & Duveau 2012; Félix et al. 2014; Ferrari et al. 2017). While there are anecdotal instances of other *Caenorhabditis* species being associated with vertebrates, including birds (Schmidt & Kuntz 1972), dogs (Kreis & Faust 1933), and humans (Scheiber 1880), no directed searches focussing on living or dead animal niches have been reported. It is therefore possible that there exists a largely undiscovered clade of vertebrate-associated *Caenorhabditis* species.

We cannot classify *C. bovis* as a “true” parasite, as opposed to an opportunistic coloniser of niches created by other pathogens, from the genome alone. Several other *Caenorhabditis* species associate with arthropods as phoretic hosts, and such phoresy is thought to serve as the major means of transport between scattered food patches (Kiontke & Sudhaus 2006). *C. bovis* is believed to be transported by dipterans (Msolla *et al*. 1989), themselves associated with parasitism of bovine ears, and may have exploited the biology of these phoretic hosts to colonise a new niche, the bovine ear. Bacterial coinfection may be a prerequisite of colonisation by *C. bovis*, may be exacerbated or encouraged by the presence of *C. bovis,* or may be initiated by *C. bovis* itself. Other species in Rhabditina offer models for this last possibility: entomopathogenic species in the genus *Heterorhabditis* carry specific bacteria which play roles in killing their arthropod larvae prey (Forst *et al*. 1997), and molluscicidal nematodes in the genus *Phasmarhabditis* induce bacterial sepsis in slugs and snail prey (Tan & Grewal 2002).

While the genome and the gene set of *C. bovis* is smaller than that of *C. elegans* and many other *Caenorhabditis* species, we identified several gene families that appear to have undergone expansion in *C. bovis.* Functional annotation of these expanded gene families revealed that several have been independently implicated in parasitism in other nematode species. P-glycoproteins (and orthologues of *pgp-11* specifically) are associated with resistance to anti-nematode drugs, and more generally in response to xenobiotic stress. FAR proteins, galectins, and serpins are actively secreted by several parasitic nematode species, and an immunomodulatory role has been proposed. It would be fascinating to explore the roles of these families (and many others) in the possible parasitic lifestyle of *C. bovis*. We note also that *C. bovis* appears to be adapted to life at 37°C in its bovine niche. This temperature is rapidly lethal to *C. elegans* (Snutch & Baillie 1983; Jones & Candido 1999) and thus *C. bovis* must have adapted to be heat resistant.

While the high-quality draft genome and the analyses presented here represent a major step forward in our understanding of this unusual and understudied *Caenorhabditis* species, it is only a beginning. Our ultimate goal is to establish long term cultures and to apply the exquisite reverse genetic toolkits available for *Caenorhabditis* to understand the biology of this species. We would like the isolates to be available to any researcher via the *Caenorhabditis* Genetics Center (CGC) and we are currently seeking the appropriate permits for export from Kenya. We hope that these cultures combined with the draft-genome sequence will enable the interrogation of the biology of *C. bovis*, including the use of CRISPR-Cas9 technology to edit or disrupt loci that might be relevant for its unusual lifestyle. It is important to note, however, that we still know very little about *C. bovis in situ*, with details of its present-day prevalence, role in bovine parasitic otitis and microbial associates remaining scarce. Therefore, any laboratory interrogation must happen alongside further study of *C. bovis* in Eastern Africa, in collaboration with local institutes and scientists.

## Methods

### Ethics statement

This study was approved by the Institutional Research Ethics Committee (IREC Reference No. 2017-08) and the Institutional Animal Care and Use Committee (IACUC Reference No. 2017-04 and 2017-04.1) at the International Livestock Research Institute, review bodies approved by the Kenyan National Commission for Science, Technology and Innovation. Approval to conduct the work was also obtained from the Department of Veterinary Services and the relevant offices of these Ministries at devolved government level. All recruited animal owners gave written, informed consent prior to their inclusion in the study.

### Sampling, nematode isolation and culture

Sampling was carried out as part of an existing surveillance programme of zoonotic disease in humans at hospitals and livestock animals at livestock markets and slaughterhouses in 3 counties of Western Kenya (Fig. 1; Table S1). A total of 44 cattle, including a range of local breeds and ages, were sampled. We restrained each animal manually and washed the external auditory canal using cotton wool soaked in physiological saline. Cotton wool samples were stored in 50 ml tubes and transported to the laboratory in a refrigerated box. We inspected 1-2 ml of saline from each sample under a dissecting microscope within 4 hours of collection. Nematodes were isolated from the saline using a pipette and placed onto nematode growth medium (NGM) or horse blood agar plates seeded with an environmentally-sourced *E. coli* strain. Plates were incubated at 37°C.

### DNA extraction

We harvested nematodes by washing each plate with phosphate-buffered saline (PBS) supplemented with 0.01% Tween20. The nematodes were washed three times with clean PBS and subsequently centrifuged to form a pellet. Pellets were stored at −40°C until extraction. To each frozen pellet, we added 600 µL of Cell Lysis Solution (Qiagen) and 20 µL of proteinase K (20 μg/μL) and incubated for four hours at 56°C. 5 µL of RNase Cocktail Enzyme Mix (Invitrogen) was subsequently added and incubated at 37°C for one hour. We added 200 µl of Protein Precipitation Solution (Qiagen) and centrifuged at 15,000 rpm for 3 minutes. The supernatant was collected in a new tube and 600 µL of isopropanol added to precipitate the DNA. We centrifuged each tube at 15,000 rpm for 3 minutes and discarded the supernatant. The resulting DNA pellets were washed twice with 70% ethanol and briefly allowed to dry before being resuspended in 100 µL of elution buffer (10 mM Tris-Cl). DNA concentration was assessed using Qubit (Thermo Scientific).

### MinION library preparation and sequencing

We sheared the DNA prior to sequencing by passing approximately 2 µg in a volume of 100 µl through a 29G insulin needle 5-10 times. Small fragments were removed using by purifying DNA with 0.5x concentration Agencourt AMPure XP beads. We followed the “one-pot” ligation protocol for preparing Oxford Nanopore SQK-LSK108 libraries (https://www.protocols.io/view/one-pot-ligation-protocol-for-oxford-nanopore-libr-k9acz2e) but with the following modifications: we added 5 µl of SQK-LSK109 adapter mix (AMX) instead of 20 µl of SQK-LSK108 AMX; we added 20 µl of NEB Ultra II Ligation Master Mix instead of 40 µl; we replaced the SQK-LSK108 adapter binding beads (ABB) with either the SQK-LSK109 long fragment buffer (LFB) or short fragment buffer (SFB). Thereafter, we followed the standard manufacturer’s instructions for preparing and loading SQK-LSK109 libraries. Libraries were loaded on to two R9.4 flow cell and run for ∼48 hours using MinKNOW version 18.12.9. Raw data metrics are presented in Table S2.

### MiSeq Sequencing

We prepared one Nextera DNA Flex library as per manufacturer’s instructions using ∼100 ng of input DNA. The library fragment size was assessed using the Agilent TapeStation and library concentration was determined using Qubit dsDNA HS Assay Kit (Thermo Scientific, USA). The library was then sequenced using the Illumina MiSeq platform with a paired-end 300 bp MiSeq reagent kit v3 (Illumina Inc, USA) at the BecA-ILRI Hub in Nairobi, Kenya.

### Genome assembly and gene prediction

Software versions and relevant parameters are available in Table S6. We base called the MinION FAST5 data using the high accuracy model in Guppy (available at https://community.nanoporetech.com). We generated a preliminary assembly using wtdbg2 (Ruan & Li 2019) and identified contaminants using taxon-annotated, GC-coverage plots (Fig. S2) as implemented in blobtools (Laetsch & Blaxter 2017a). Reads were mapped to the preliminary assembly using minimap2 (Li 2018) and the likely taxonomic origin of each contig was determined by searching NCBI nucleotide ‘nt’ or UniProt Reference Proteomes (Pundir *et al*. 2017) using NCBI-BLAST+ (Camacho *et al*. 2009) or DIAMOND (Buchfink *et al*. 2015), respectively. Reads originating from contaminant organisms were discarded. We generated the final assembly using wtdbg2. Sequencing errors were initially corrected by aligning the MinION reads to the assembly using minimap2 and performing four iterations of Racon (Vaser *et al*. 2017) followed by a single iteration of Medaka (available at https://github.com/nanoporetech/medaka). Any remaining errors were corrected by aligning the Illumina MiSeq reads to the assembly using BWA-MEM (Li 2013) and performing two iterations of Racon followed by two iterations of Pilon (Walker *et al*. 2014).

Repeat sequences were identified *de novo* using RepeatModeler (Smit & Hubley 2010) and subsequently masked using RepeatMasker (Smit *et al*. 1996). Protein-coding genes were predicted using BRAKER (Hoff *et al*. 2016), using proteins sequences from nematode-specific EggNOG database (which comprises sequences from *C. elegans*, *C. briggsae*, *C. remanei*, *C. japonica, Pristionchus pacificus* and *Trichinella spiralis*) (Huerta-Cepas *et al*. 2016) as homology evidence. Genome assembly and gene set completeness were assessed using BUSCO with the ‘nematoda_odb9’ database (Simão *et al*. 2015). We used Jellyfish (Marçais & Kingsford 2011) to count kmers in the Illumina MiSeq reads and used GenomeScope (Vurture *et al*. 2017) to estimate genome size and heterozygosity.

To assign *C. bovis* contigs to chromosomes, we identified one-to-one orthologues between *C. bovis* and *C. elegans* using a reciprocal best BLAST hit approach. Both proteomes were filtered so that they contained only the longest-isoform per gene and searched against each other using BLASTp. Protein pairs which had reciprocal best BLAST hits with e-values < 1e-25 and a query coverage >75% were declared as one-to-one orthologues. *C. bovis* contigs containing 10 or more *C. elegans* orthologues were assigned to the chromosome containing the majority of the *C. elegans* orthologues.

### Orthology clustering and phylogenetic inference

Accession details for all data used in this analysis are available in Table S7. We selected the protein sequence of the longest isoform of each protein-coding gene in *C. bovis*, 32 other species of *Caenorhabditis,* and the two outgroup taxa, *Diploscapter coronatus* and *Diploscapter pachys.* OrthoFinder (Emms & Kelly 2015) was used to cluster all protein sequences into putatively orthologous groups (OGs) using the default inflation value of 1.5. OGs containing loci which were present in at least 75% of species and which were, on average, single copy (mean count per species < 1.3) were selected. We aligned each selected OG using MAFFT (Katoh & Standley 2013) and generated a maximum likelihood tree along with 1000 ultrafast bootstraps (Hoang *et al*. 2018) using IQ-TREE (Nguyen *et al*. 2015), allowing the best-fitting substitution model to be selected automatically (Kalyaanamoorthy *et al*. 2017). Each tree was screened by PhyloTreePruner (Kocot *et al*. 2013), collapsing nodes with bootstrap support <90, and any OGs containing paralogues were discarded. If two representative sequences were present for any species (i.e., “in-paralogues”) after this paralogue screening step, only the longest of the two sequences was retained. We then realigned the remaining OGs using MAFFT and trimmed spuriously aligned regions using trimAl (Capella-Gutiérrez *et al*. 2009). The trimmed alignments were subsequently concatenated to form a supermatrix using catfasta2phyml (available at https://github.com/nylander/catfasta2phyml). We inferred the species tree using IQ-TREE with the general time reversible model (GTR) with gamma-distributed rate variation among sites. The resulting tree was visualized using the iTOL web server (Letunic & Bork 2016).

### Identification of expanded/contracted gene families

We used KinFin (Laetsch & Blaxter 2017b) to identify gene families that were expanded or reduced in *C. bovis*. We searched the longest isoform of each protein-coding gene in all 35 species against the Pfam and SignalP databases using InterProScan (Jones *et al*. 2014) and provided the resulting annotations to KinFin. Expanded and contracted gene families were identified using log2-transformed ratios of counts in *C. bovis* to mean or median counts in other species (Table S3). Gene trees were inferred using IQ-TREE as described previously.

## Acknowledgements

We thank the University of Edinburgh’s Davis Fund for the generous support of this project. We thank members of ILRI ZooLink team for their assistance and helpful discussions. We thank Karin Kiontke and David Fitch for assistance with identification of isolated nematodes. This work was supported by the Biotechnology and Biological Sciences Research Council, the Department for International Development, the Economic & Social Research Council, the Medical Research Council, the Natural Environment Research Council and the Defence Science & Technology Laboratory, under the Zoonoses and Emerging Livestock Systems (ZELS) programme, grant reference BB/L019019/1. It also received support from the CGIAR Research Program on Agriculture for Nutrition and Health (A4NH), led by the International Food Policy Research Institute (IFPRI). We also acknowledge the CGIAR Fund Donors (http://www.cgiar.org/funders/). LS is funded by a Baillie Gifford Studentship. SR is funded by MRC Grant Ref. MR/N013166/1

## Author contributions

LS, MB and EMF conceptualised the study. LS, LCF, MKM, KM, COO, AO and JO performed field sampling. LS performed nematode isolation. LS, SR and SMN performed nematode culture. LS and SR performed DNA extraction and MinION sequencing. EM performed MiSeq sequencing. LS performed genome assembly, phylogenetic analysis, and comparative genomic analysis. LS and MB wrote the manuscript. All authors commented on and approved the final version of the manuscript.

## Data availability

Raw sequence data and the genome assembly have been deposited in the relevant INSDC databases under the accession PRJEB34497. The assembly and gene set are available to browse, query, and download at http://www.caenorhabditis.org. Data files associated with this study have been deposited in Zenodo under the accession 10.5281/zenodo.3437796.

## Supplementary Information

**Table S1:** Sampling locations and number of animals sampled

| **Site** | **Name** | **GPS coordinates** | **Type** | **Date** | **Number of animals sampled** |
| --- | --- | --- | --- | --- | --- |
| 1 | Amukura | 0.55662731, 34.26680546 | SH | 26/03/2019 | 3 |
| 2 | Kimilli | 0.78995062, 34.72008678 | SH | 28/03/2019 | 7 |
| 3 | Ikolomani | 0.203020438, 34.66886937 | LM | 02/04/2019 | 7 |
| 4 | Myanga | 0.561452038, 34.38950072 | LM | 03/04/2019 | 7 |
| 5 | Angurai | 0.637058568, 34.2727974 | LM | 03/04/2019 | 7 |
| 6 | Malaba | 0.63711942, 34.27281931 | SH | 03/04/2019 | 3 |
| 7 | Chwele | 0.739193074, 34.58697939 | LM | 08/04/2019 | 7 |
| 8 | Chwele | 0.73913342, 34.58692581 | SH | 08/04/2019 | 2 |

**LM:** Livestock market, **SH:** slaughterhouse. *C. bovis* was isolated from an individual sampled at a livestock market in Chwele (site 7).

**Table S2:** MinION sequencing statistics

|  | ***Run 1*** | ***Run 2*** |
| --- | --- | --- |
| Total data (Gbp) | 4.42 | 6.83 |
| Read count | 955,868 | 4,857,912 |
| Read length N50 (kbp) | 11.41 | 4.32<sup>§</sup> |
| Mean read length (kbp) | 4.62 | 1.41 |
| Longest read (kbp) | 242.02 | 172.39 |

<sup>§</sup>Library prepared using the SQK-LS109 short fragment buffer (Oxford Nanopore) as the DNA appeared to be highly fragmented.

**Table S3:** Gene families that are expanded/contracted in *C. bovis* relative to 32 other *Caenorhabditis* species (Too large to display in text; available as separate .xls file)

**Table S4:** Repetitive content in the C*. bovis* and *C. elegans* genomes

|  | ***C. bovis*** | ***C. elegans*** |
| --- | --- | --- |
| Total repeat content (Mb)<sup>§</sup> | 8.60 (13.7%) | 18.1 (18.1%) |
| SINEs (Mb) | 0.03 (0.05%) | 0.19 (0.19%) |
| LINEs (Mb) | 0.17 (0.27%) | 0.38 (0.38%) |
| LTR elements (Mb) | 0.35 (0.56%) | 0.20 (0.20%) |
| DNA elements (Mb) | 2.58 (4.12%) | 8.89 (8.86%) |
| Unclassified (Mb) | 4.32 (6.88%) | 4.22 (4.21%) |

<sup>§</sup>Includes simple/low complexity repeats, satellites and small RNA in addition to transposable elements. Repeat content estimated *de novo* using RepeatModeller and RepeatMasker.

**Table S5:** Gene structure statistics^§^ of *C. bovis* and *C. elegans*

|  | All genes |  | Orthologues only |  |
| --- | --- | --- | --- | --- |
|  | *C. bovis* | *C. elegans* | *C. bovis* | *C. elegans* |
| gene count | 13128 | 20208 | 7706 | 7706 |
| total span of genes (excluding UTRs) (bp) | 35062444 | 57199834 | 21555520 | 30210405 |
| total exon span (bp) | 18715784 | 24639393 | 11766915 | 11906370 |
| mean exon span per gene (bp) | 1426 | 1219 | 1527 | 1545 |
| total exon count | 117212 | 122373 | 73985 | 58546 |
| mean exon count per gene | 8.93 | 6.06 | 9.60 | 7.60 |
| total intron span (bp) | 16346660 | 32560441 | 9788605 | 18304035 |
| mean intron span per gene (bp) | 1245 | 1611 | 1270 | 2375 |
| total intron count | 104084 | 102165 | 66279 | 50840 |
| mean intron count per gene | 7.93 | 5.06 | 8.60 | 6.60 |
| mean intron length (bp) | 157 | 319 | 148 | 360 |

<sup>§</sup>Counts shown are for the longest isoform of each gene. Untranslated regions (UTRs) were not annotated for *C. bovis* and so were not considered in either species. WormBase ParaSite version WBPS12 of the *C. elegans* genome was used.

**Table S6:**
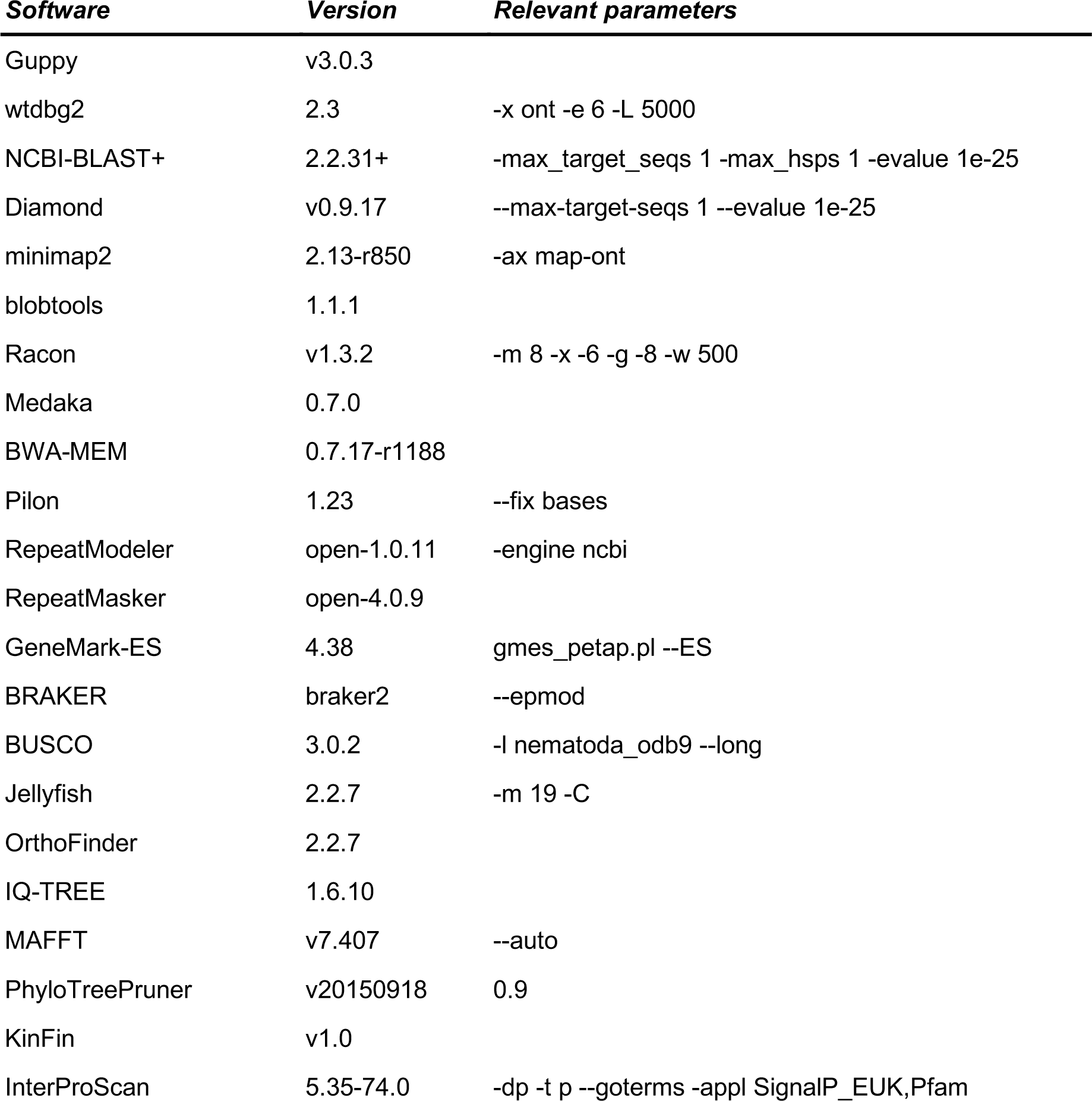
Software versions and relevant parameters

**Table S7:**
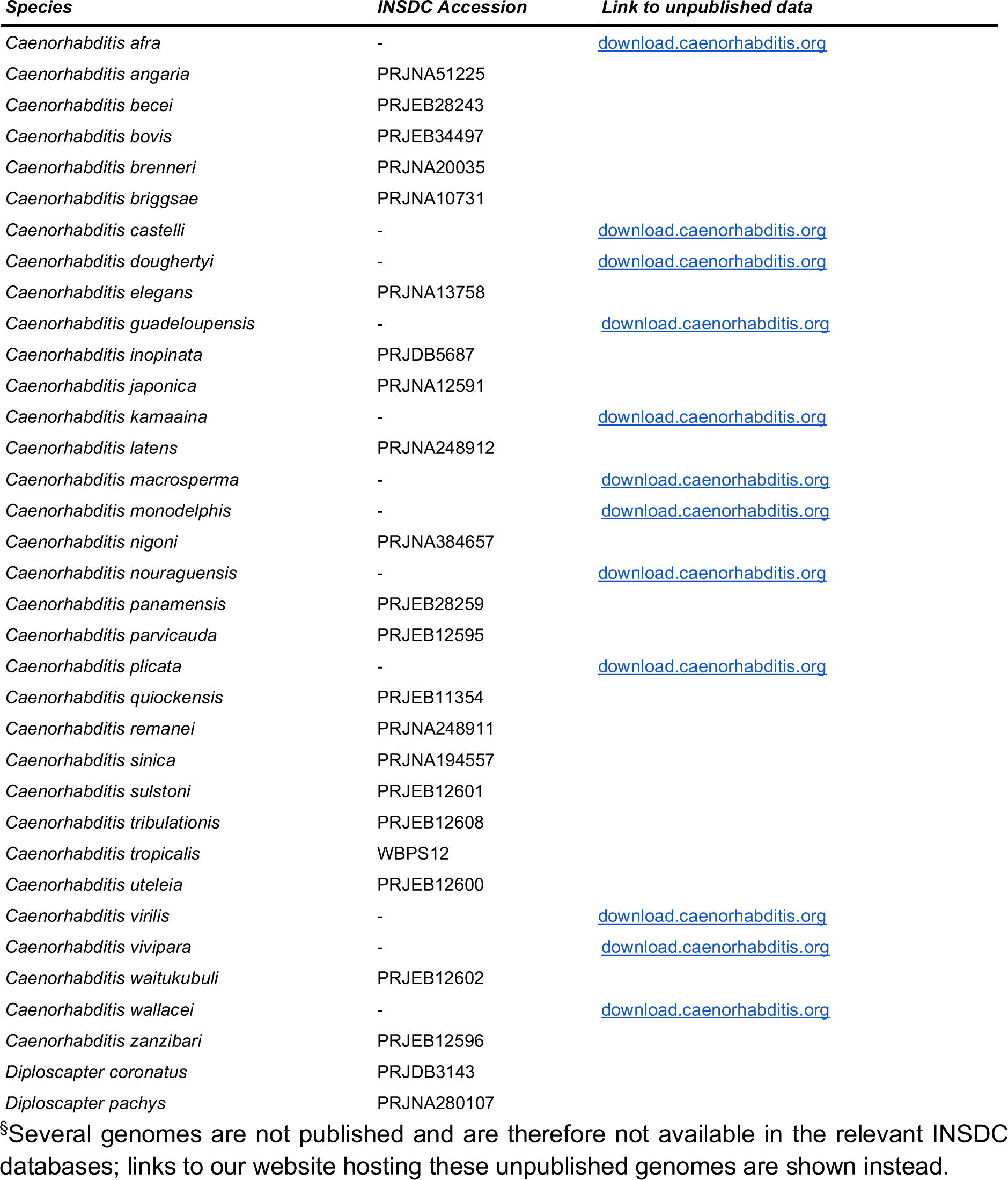
Accessions^§^ to all data used in orthology clustering and phylogenetic analysis

**Figure S1:**
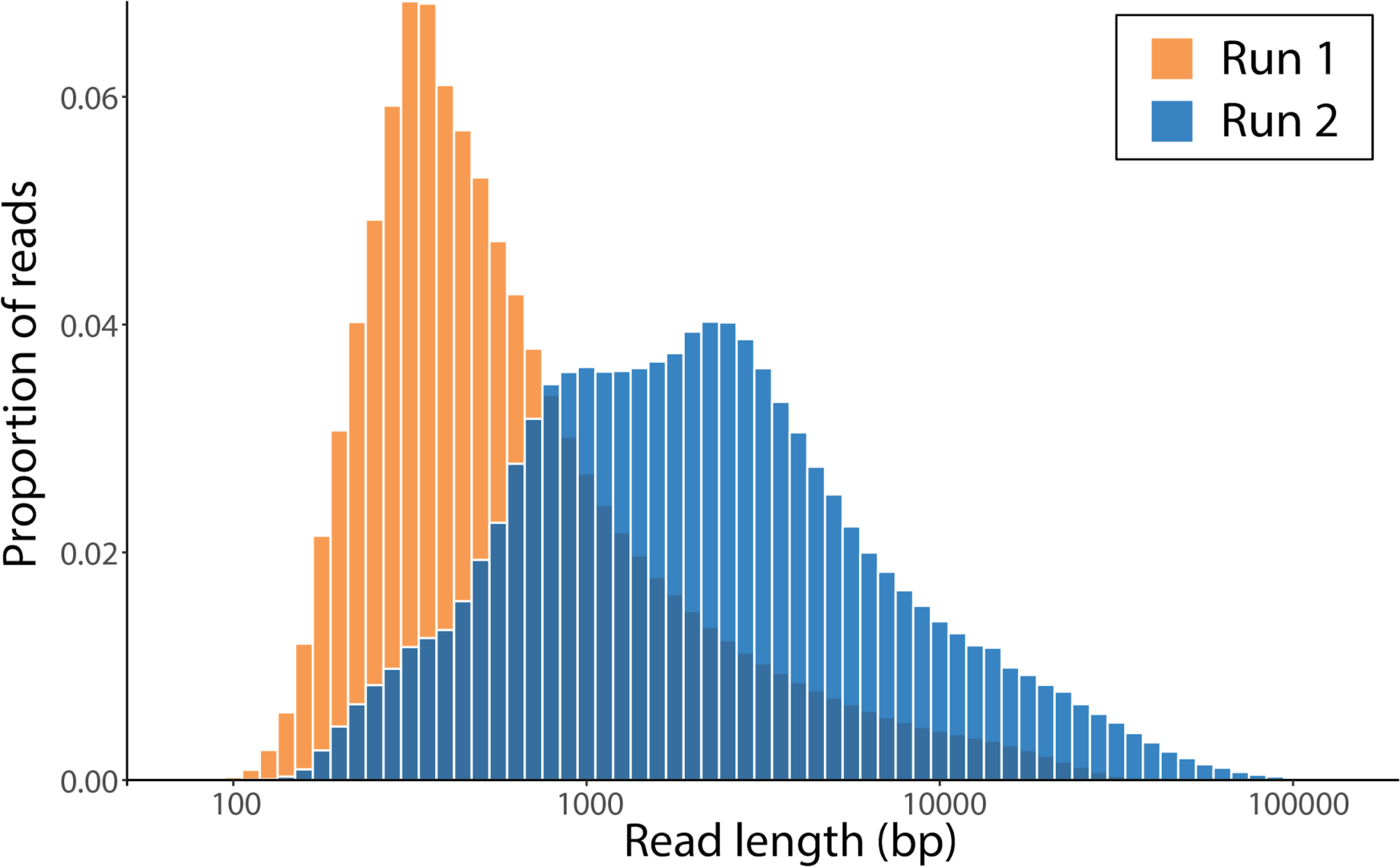
MinION read length histogram. For Run 2, we prepared the library using the SQK-LS109 short fragment buffer (Oxford Nanopore) as the DNA appeared to be highly fragmented.

**Figure S2:**
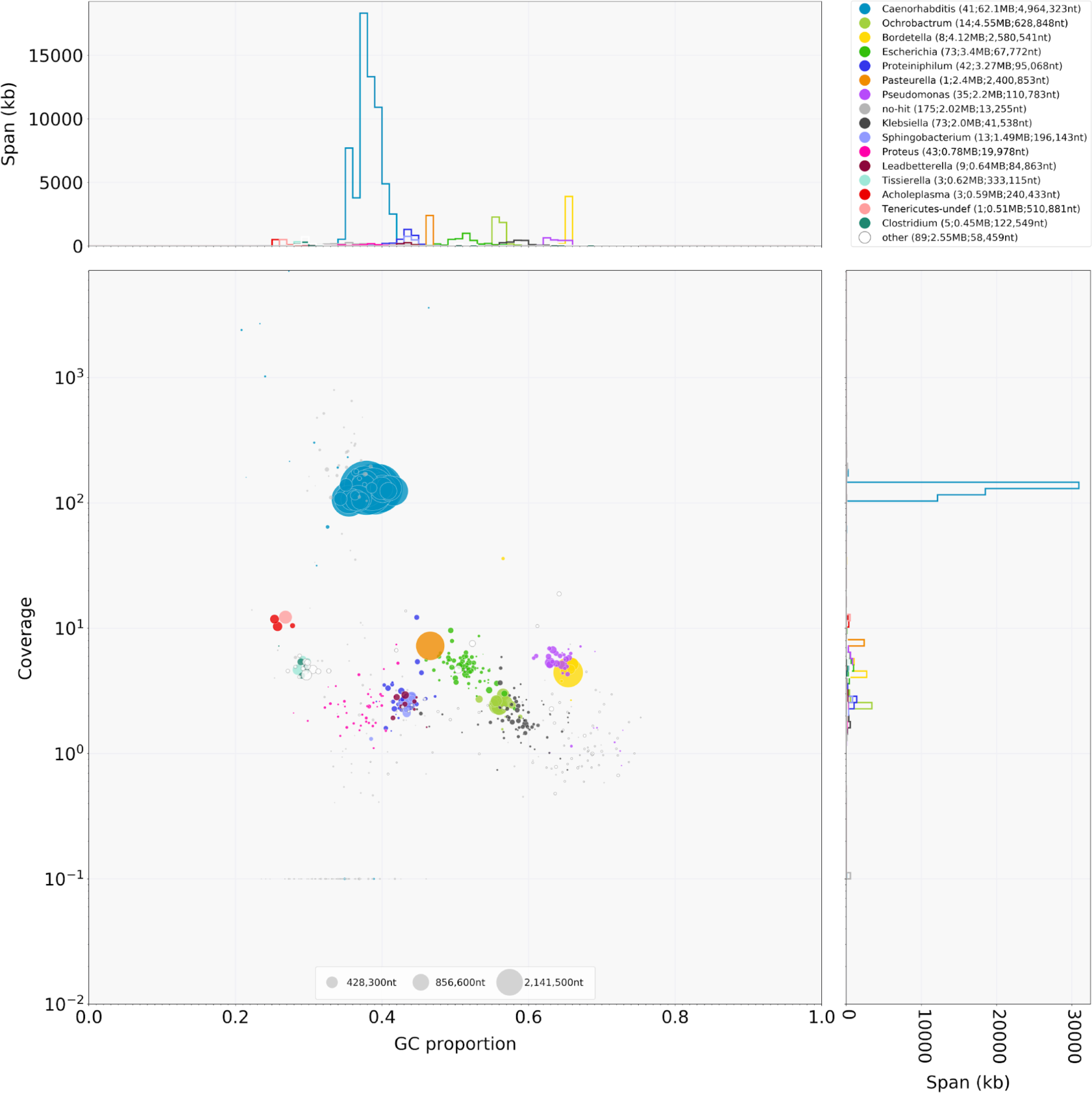
Taxon-annotation GC-coverage plot of a preliminary assembly of the *C. bovis* genome. Results of NCBI-BLAST+ and Diamond searches of NCBI nucleotide ‘nt’ or UniProt Reference Proteomes databases were provided to blobtools which assigned taxonomy (using the ‘bestsumorder’ taxonomy rule). Coverage of each contig in the MinION read set is shown. Several contigs have top hits to bacterial species which are known mammalian pathogens, including *Pasteurella multocida* (cause of haemorrhagic septicaemia in cattle (Annas *et al.* 2014)), *Ochrobactrum anthropi,* and *Bordetella petrii* (both of which have been associated with opportunistic infections in humans (Zelazny *et al.* 2013; Aguilera-Arreola *et al.* 2018)).

**Figure S3:**
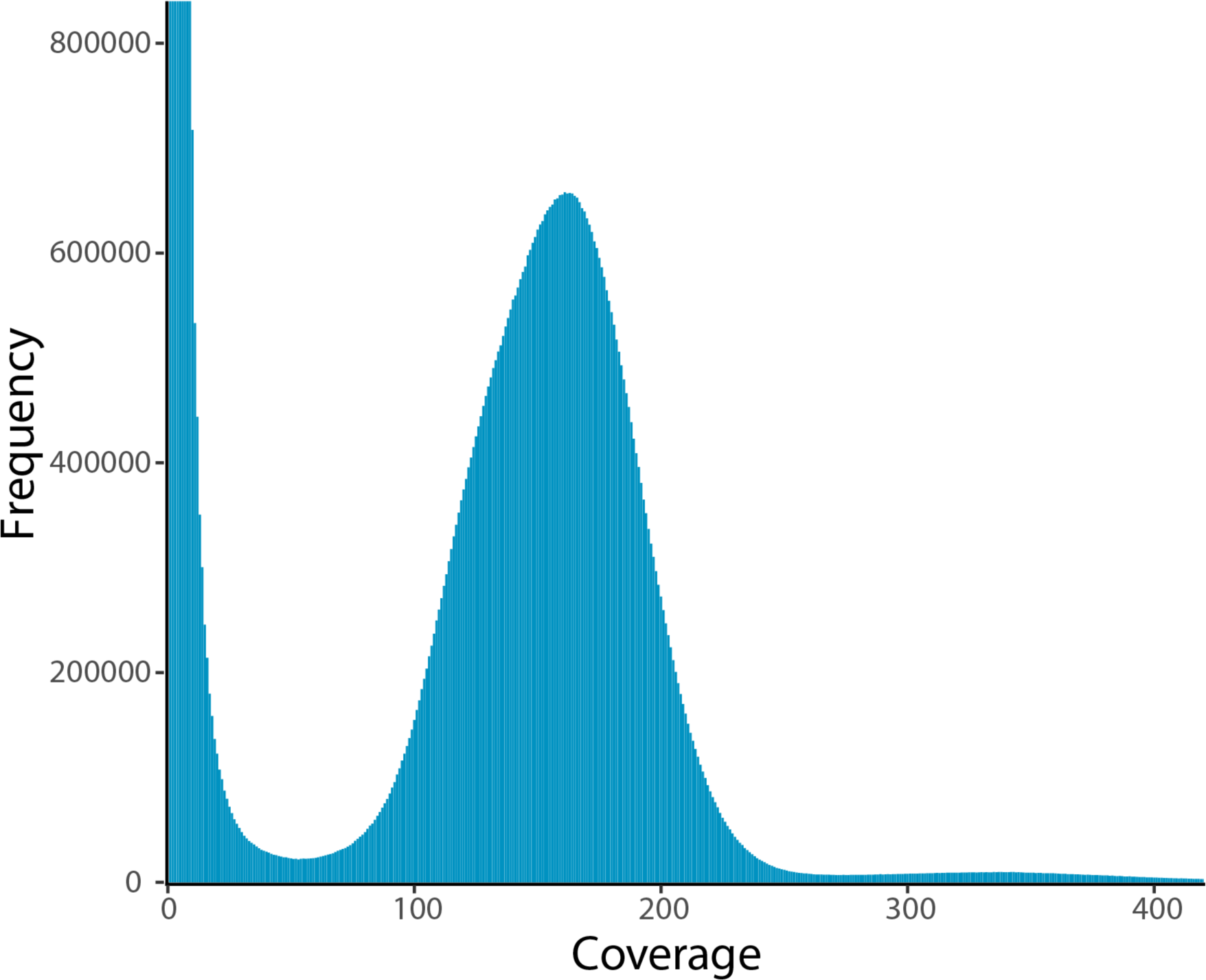
Kmer spectra of Illumina short-reads. Kmers of length 19 were counted in Illumina MiSeq reads using Jellyfish. GenomeScope estimated 0.126% heterozygosity and a genome size of 61.3 Mb.

**Figure S4:**
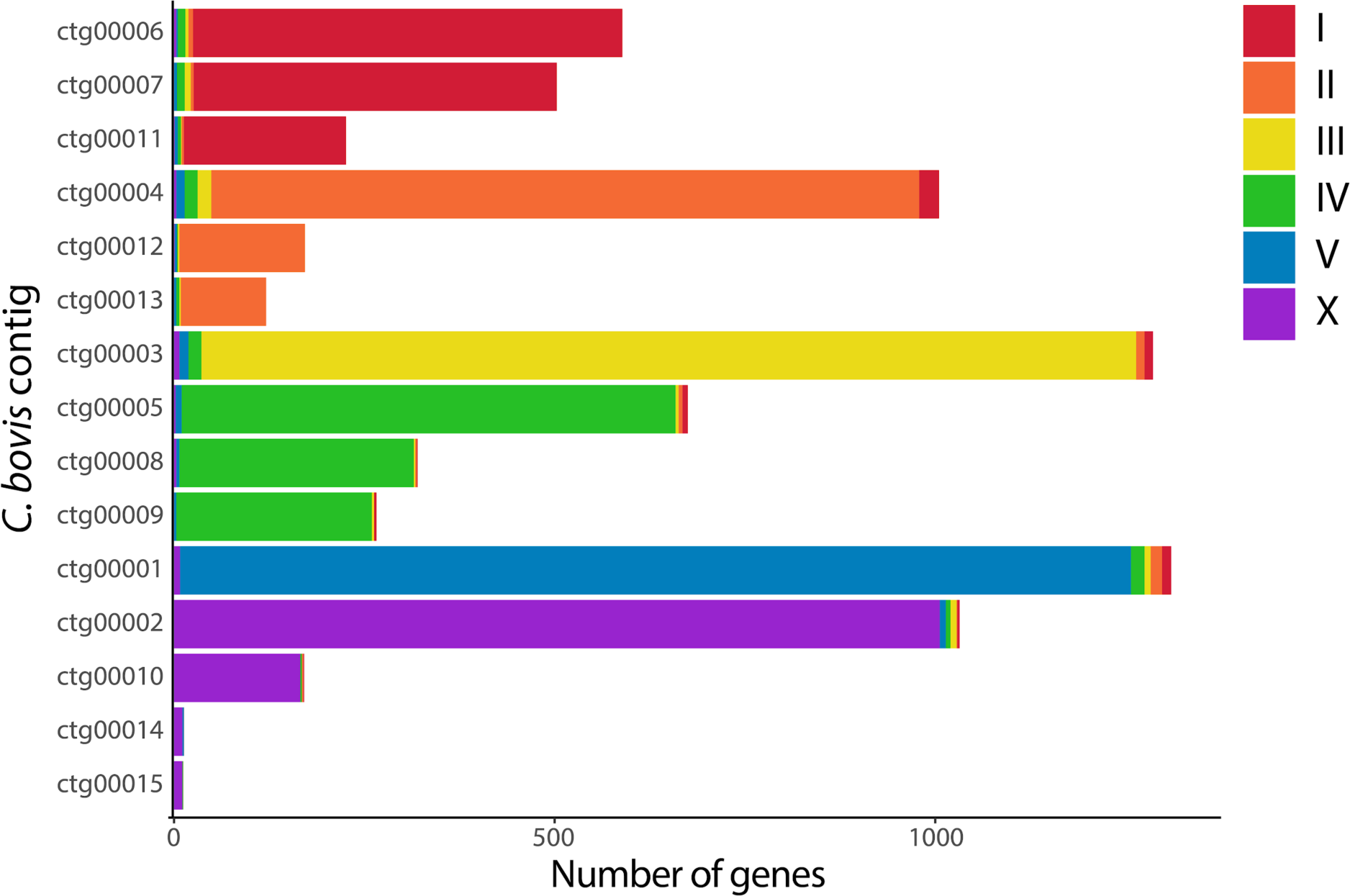
Composition of 15 *C. bovis* contigs. 15 contigs (representing 99.4% of the *C. bovis* assembly) are shown. Bars represent the number of *C. bovis* genes with a *C. elegans* orthologue on each contig; bars are coloured by the chromosome location of the *C. elegans* orthologue. 20 contigs, each containing fewer than 10 *C. elegans* orthologues and cumulatively spanning 0.39 Mb, are not shown.

**Figure S5:**
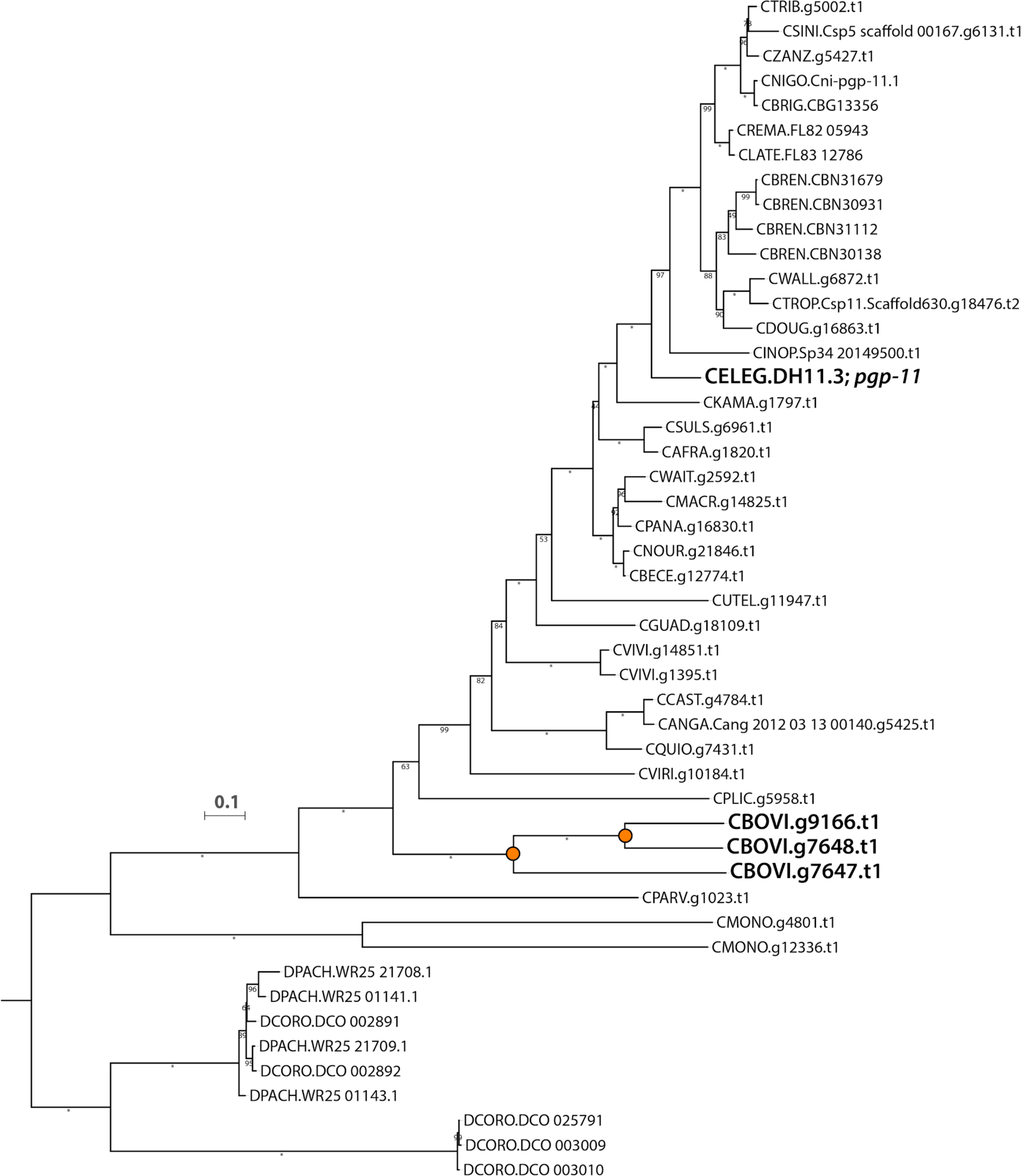
Two duplications of *pgp-11 in C. bovis.* Maximum likelihood gene tree of the orthogroup containing *pgp-11* (OG0002589) inferred using the LG+I+G4 substitution model. *C. bovis* duplications are indicated by orange circles on internal nodes. Branch lengths represent the number of substitutions per site; scale is shown. Bootstrap values are displayed as branch annotations; “*” = 100.

**Figure S6:**
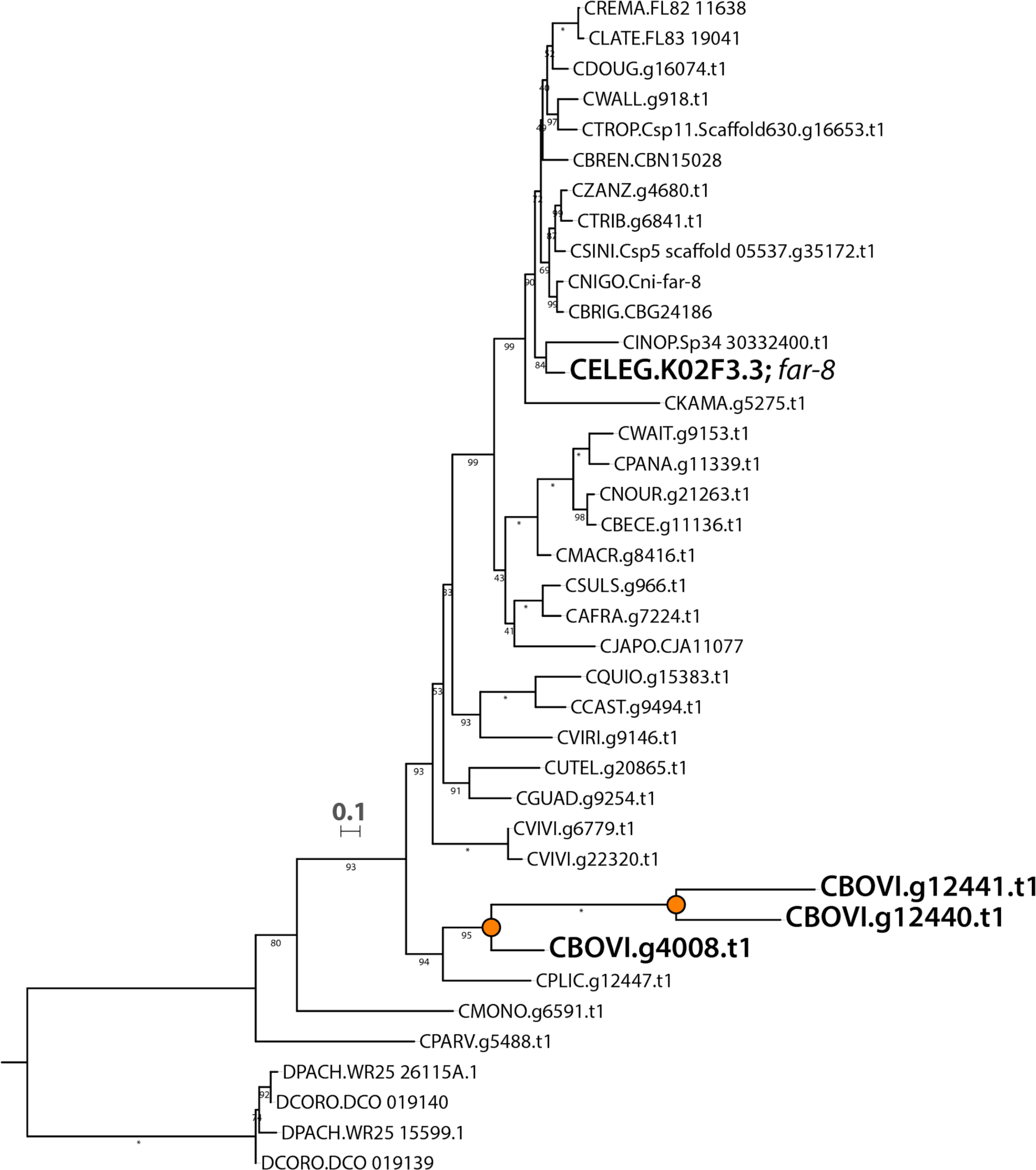
Two duplications of *far-8* in *C. bovis*. Maximum likelihood gene tree of the orthogroup containing *far-8* (OG0006033) inferred using the LG+G4 substitution model. *C. bovis* duplications are indicated by orange circles on internal nodes. Branch lengths represent the number of substitutions per site; scale is shown. Bootstrap values are displayed as branch annotations; “*” = 100.

**Figure S7:**
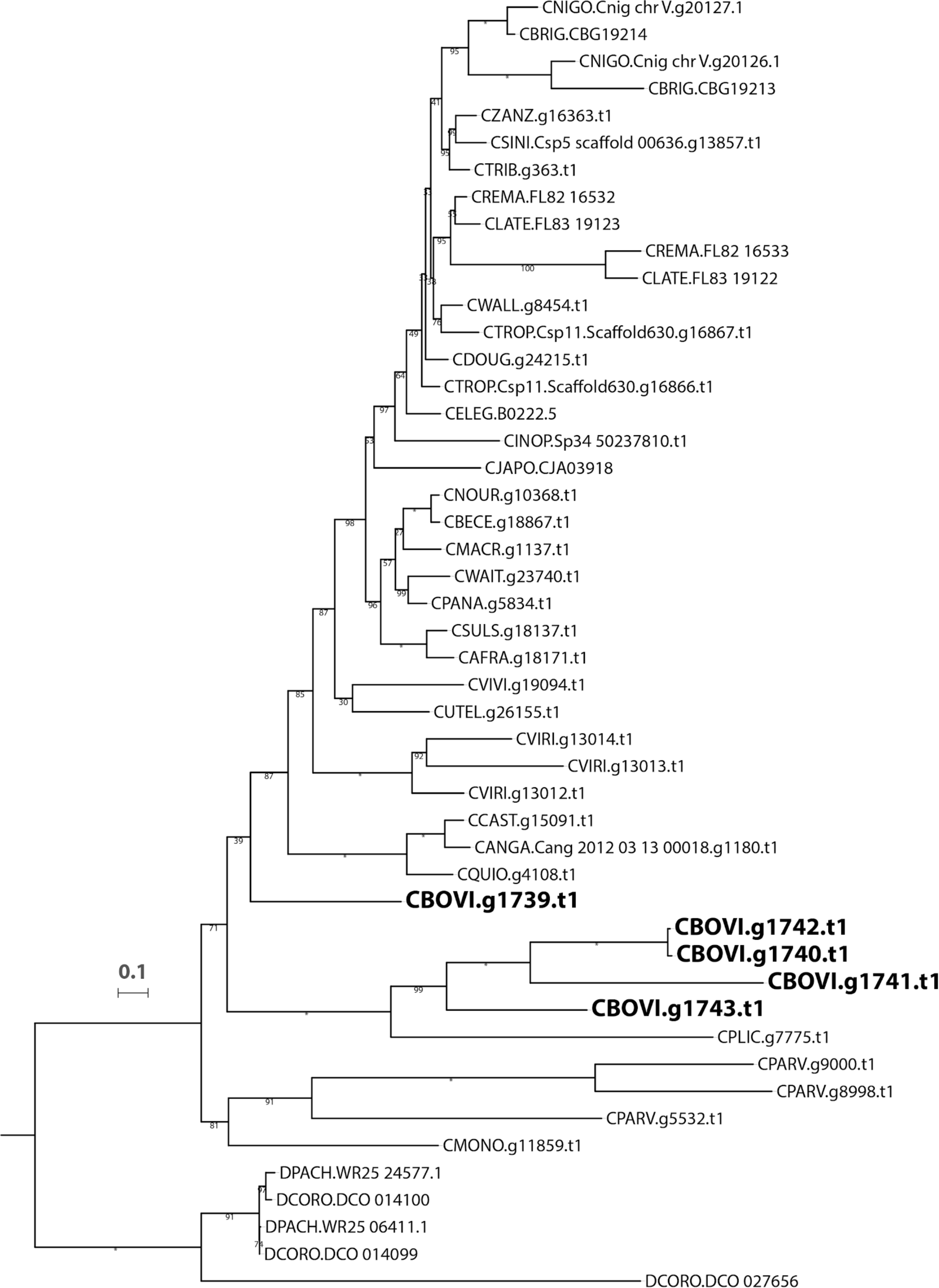
Expansion of Kunitz-type serine protease inhibitor domain containing proteins in *C. bovis*. Maximum likelihood gene tree of the orthogroup containing a family of serine protease inhibitors (OG0002693) inferred using the WAG+I+G4 substitution model. Branch lengths represent the number of substitutions per site; scale is shown. Bootstrap values are displayed as branch annotations; “*” = 100.

