## Supplementary figures and images for "The genome of *Caenorhabditis bovis*"

### Figure S1

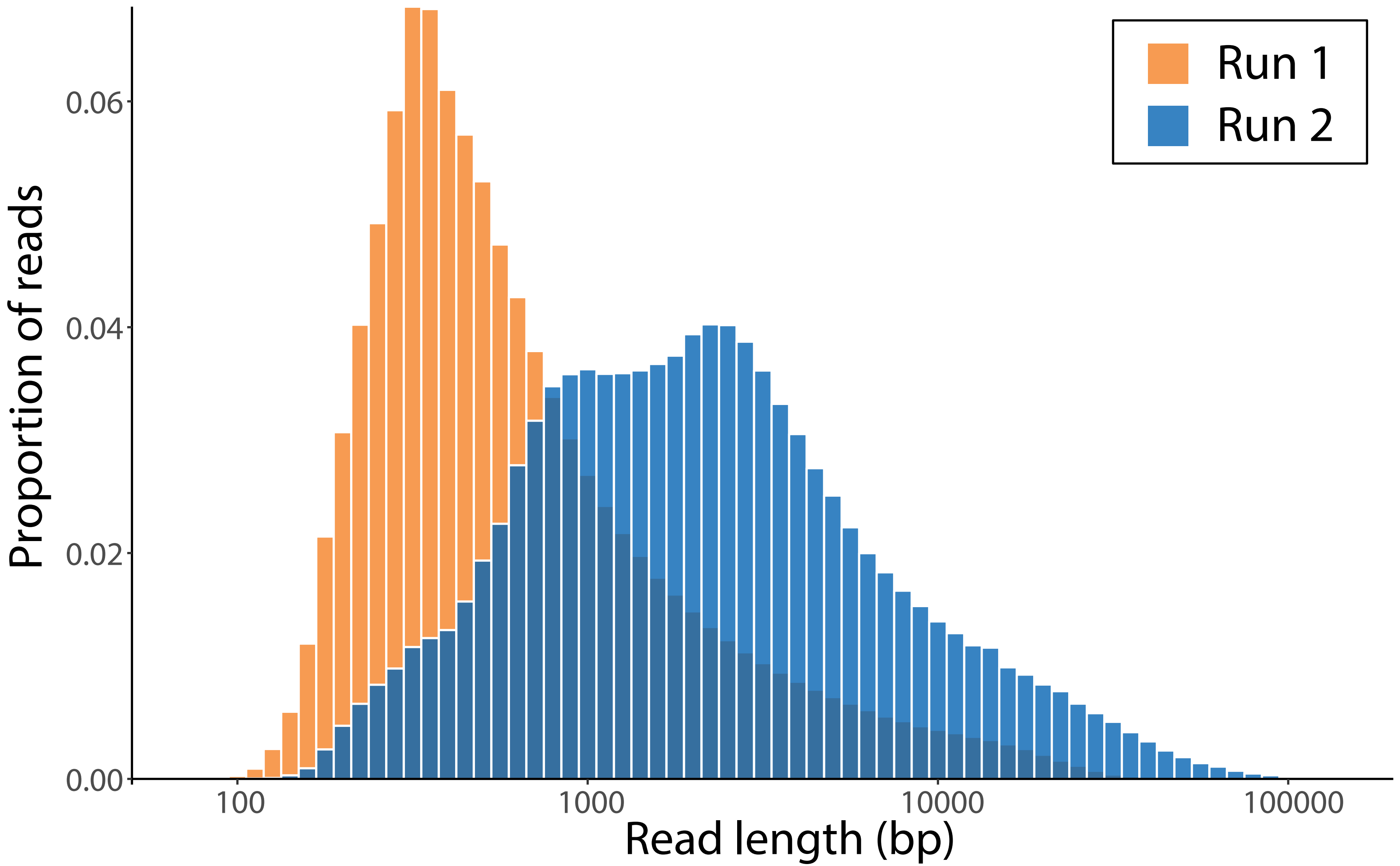

### Figure S2

CBOVI.blobDB.json.bestsumorder.genus.p16.span.1000.blobplot.covsum

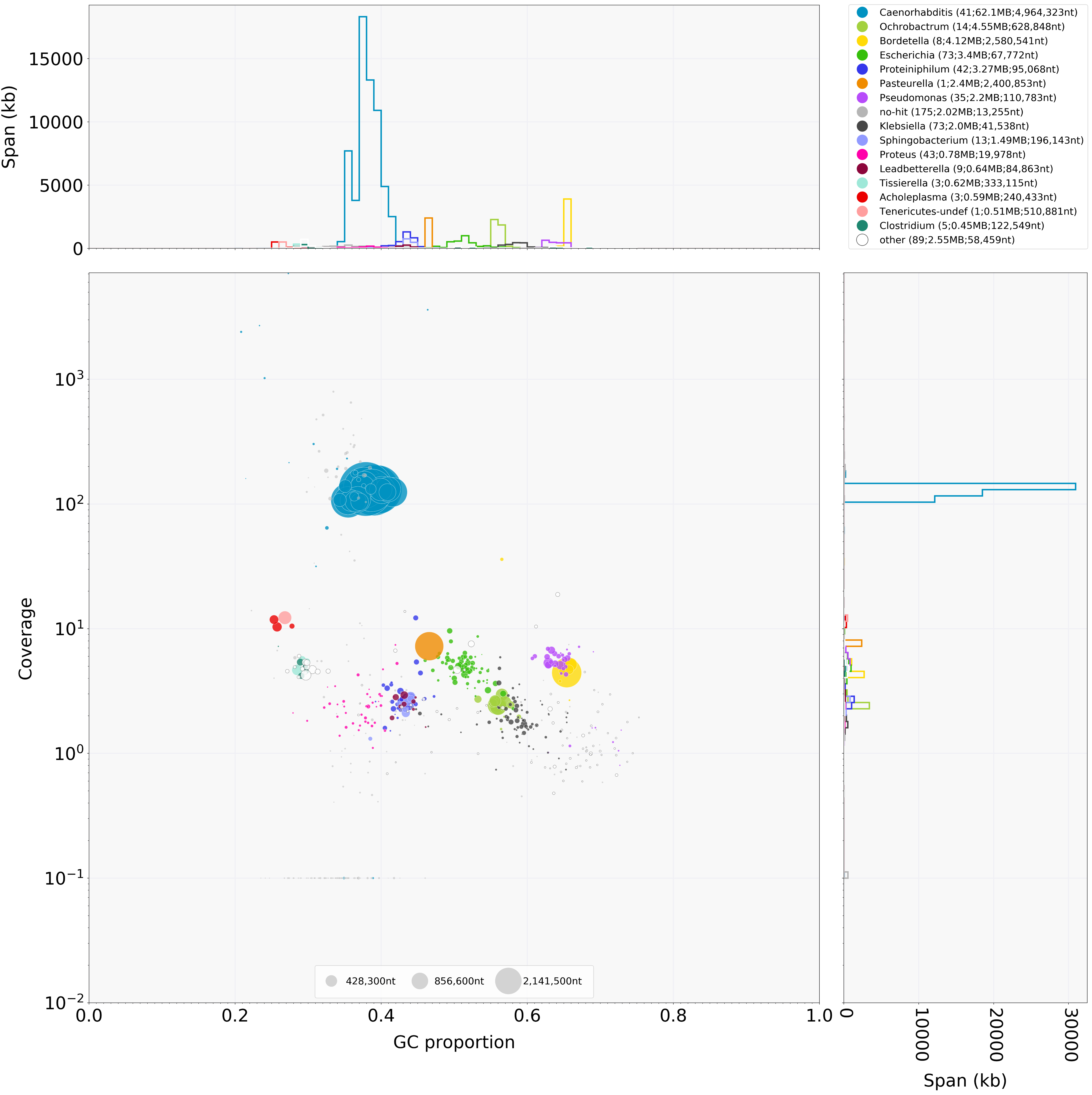

### Figure S3

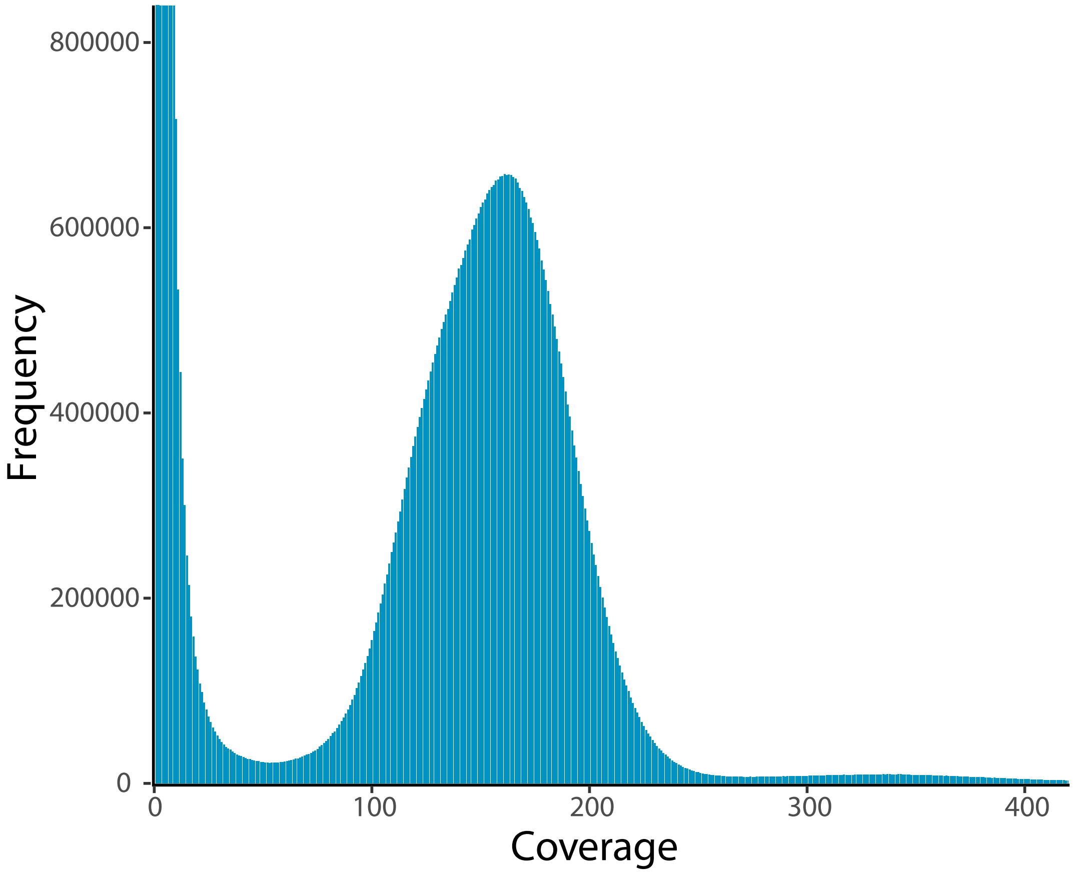

### Figure S4

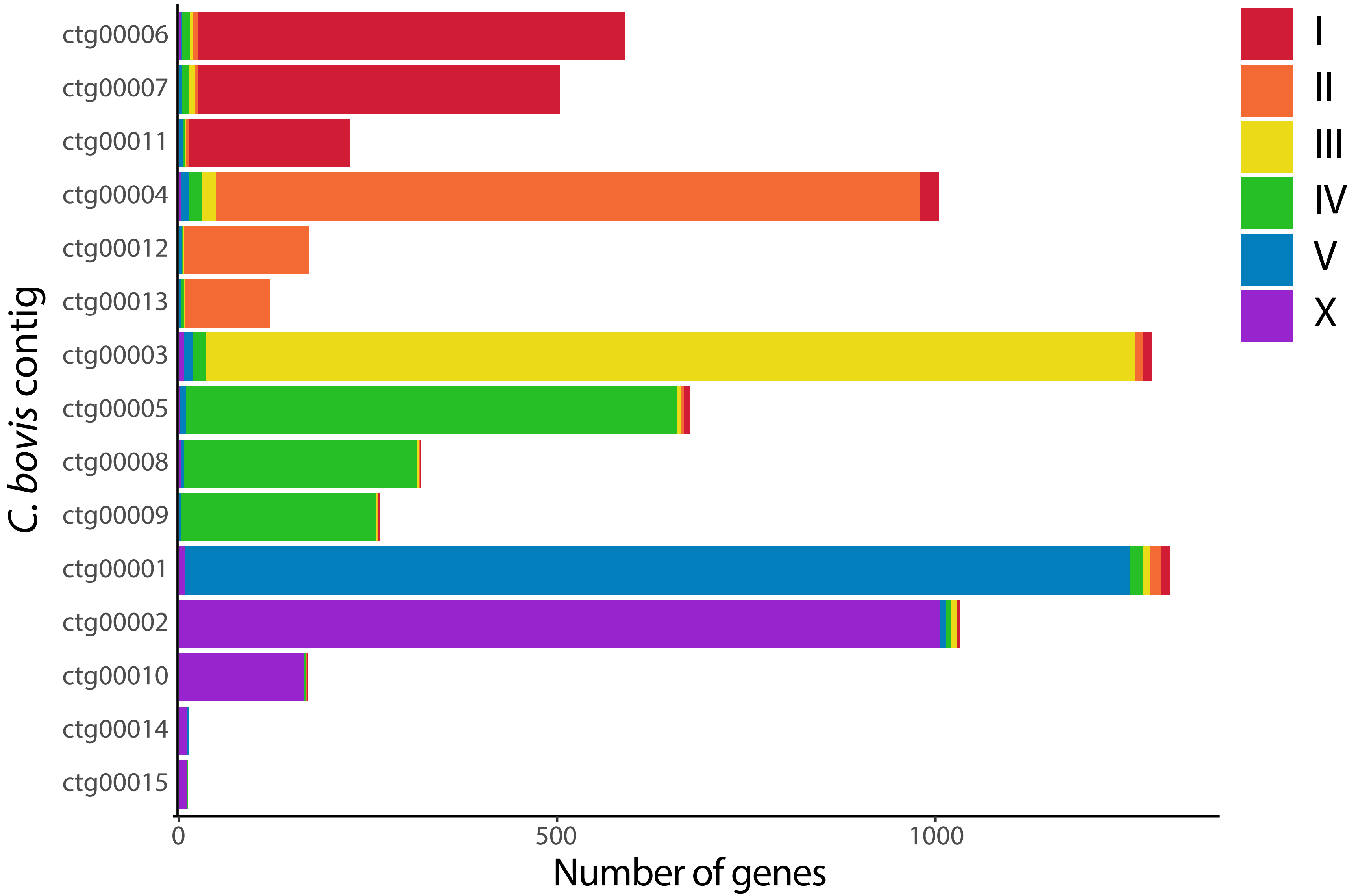

### Figure S5

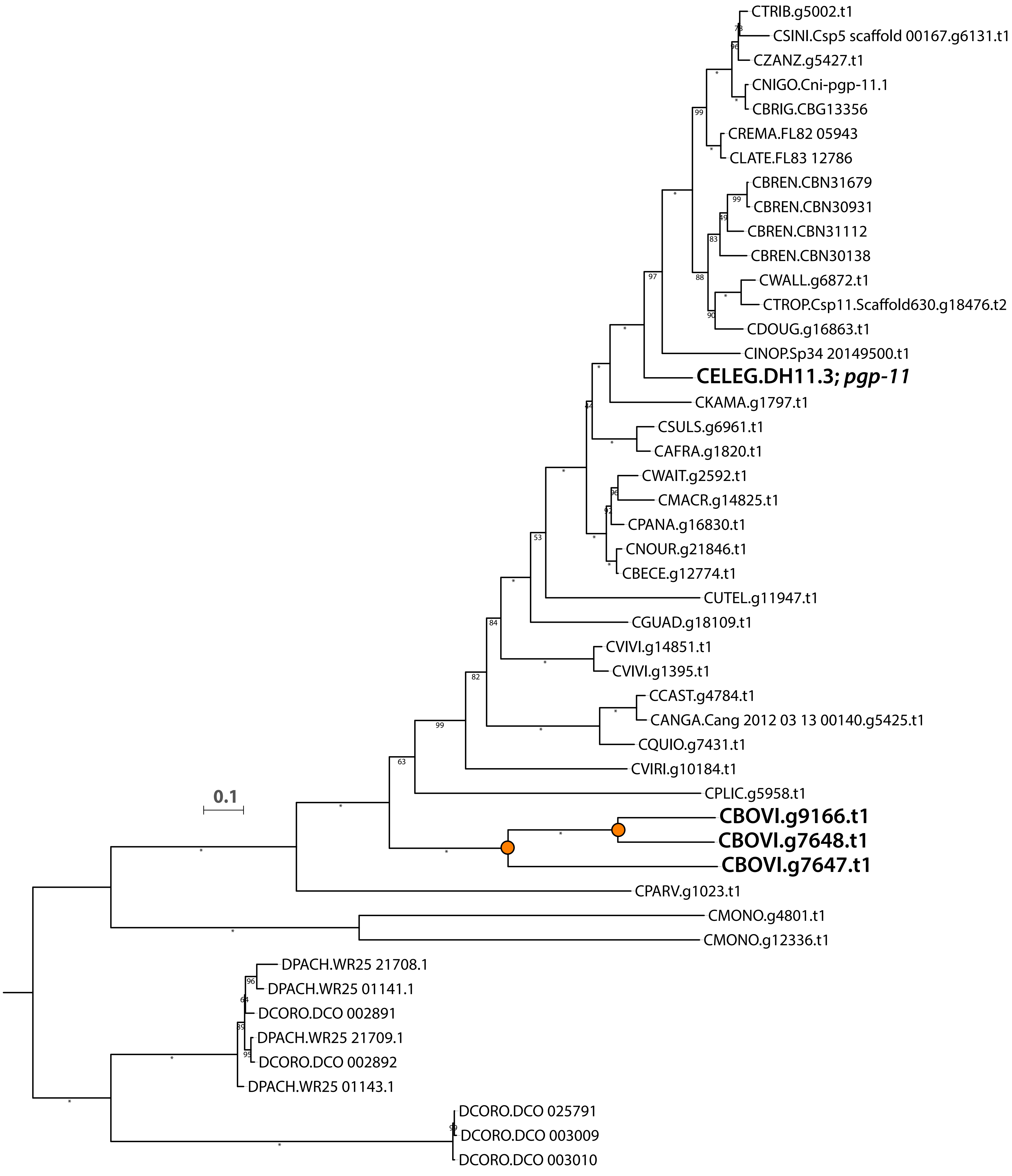

### Figure S6

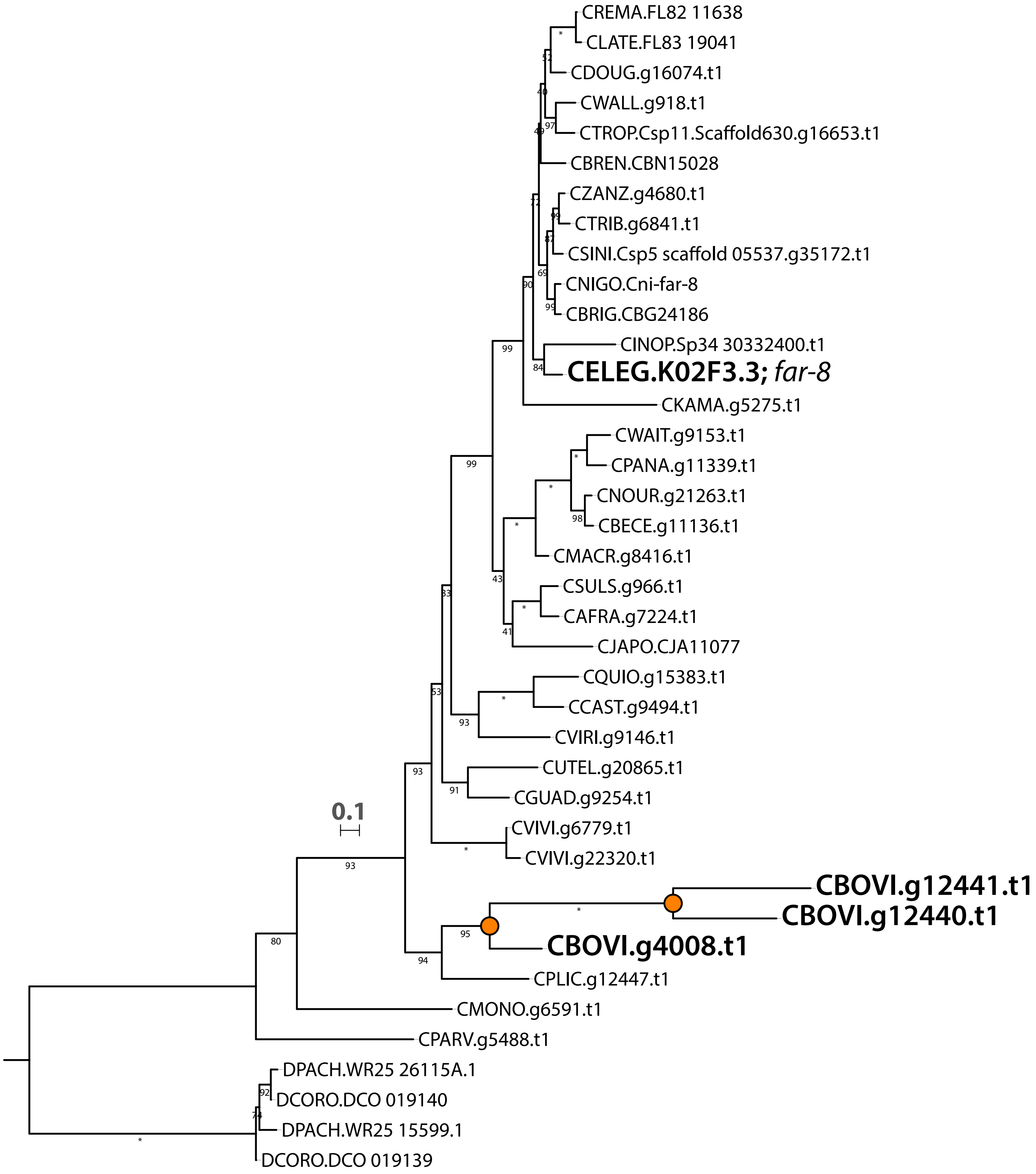

### Figure S7

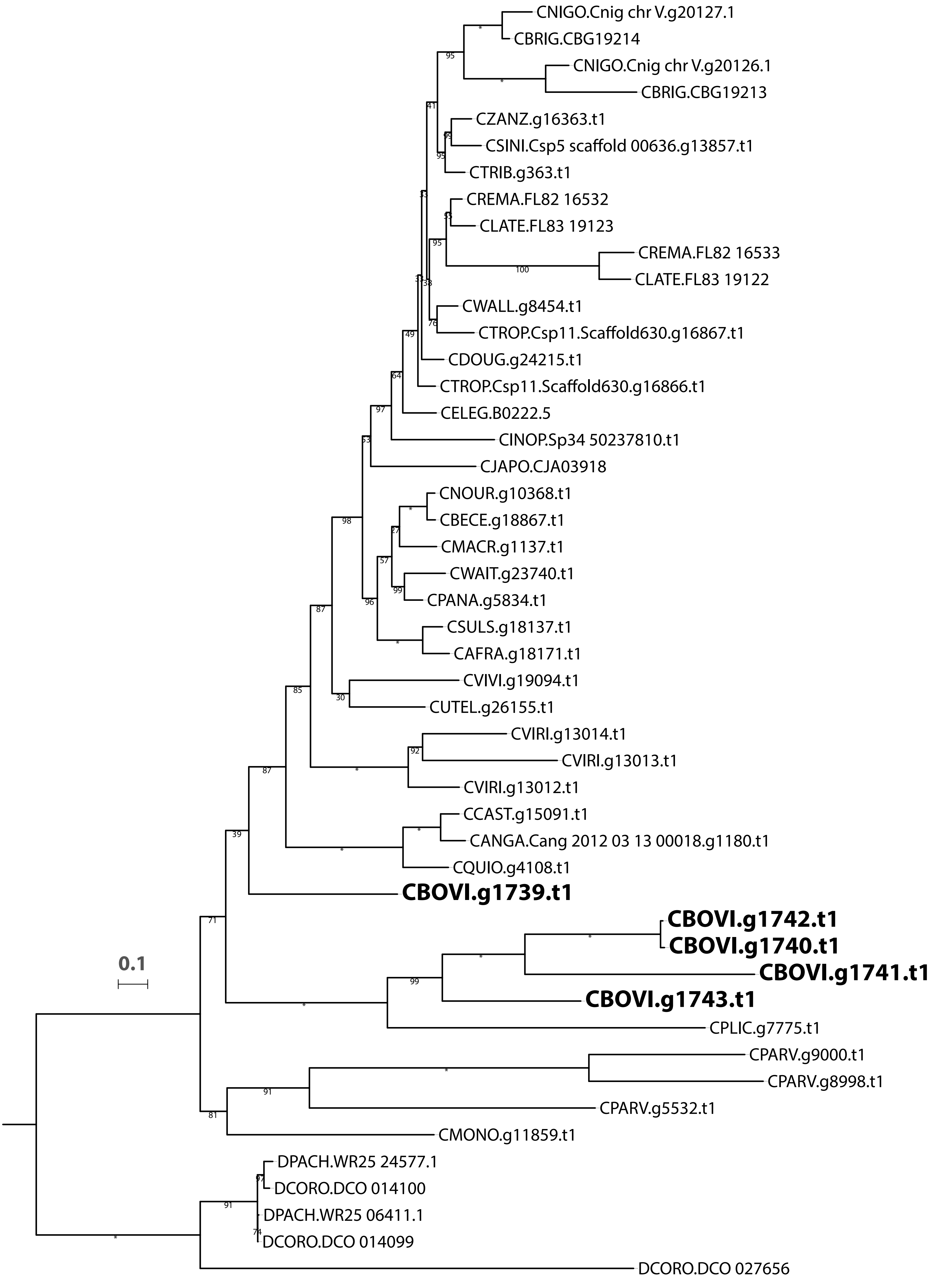
